# Specific inhibition and disinhibition in the higher-order structure of a cortical connectome

**DOI:** 10.1101/2023.12.22.573036

**Authors:** Michael W. Reimann, Daniela Egas Santander, András Ecker, Eilif B. Muller

## Abstract

Neurons are thought to act as parts of assemblies with strong internal excitatory connectivity. Conversely, inhibition is often reduced to blanket inhibition with no targeting specificity. We analyzed the structure of excitation and inhibition in the MICrONS *mm*^3^ dataset, an electron microscopic reconstruction of a piece of cortical tissue. We found that excitation was structured around a feed-forward flow in large non-random neuron motifs with a structure of information flow from a small number of sources to a larger number of potential targets. Inhibitory neurons connected with neurons in specific sequential positions of these motifs, implementing targeted and symmetrical competition between them. None of these trends are detectable in only pairwise connectivity, demonstrating that inhibition is structured by these large motifs. While descriptions of inhibition in cortical circuits range from non-specific blanket-inhibition to targeted, our results describe a form of targeting specificity existing in the higher-order structure of the connectome. These findings have important implications for the role of inhibition in learning and synaptic plasticity.

## INTRODUCTION

Assemblies are groups of neurons that tend to fire together, and that have been observed in both hippocampal and cortical activity (Hebb, 1949; Harris et al., 2003; Dragoi and Buzsáki, 2006; Carrillo-Reid et al., 2015). Similarly, in simulation studies neurons are often wired into clusters that produce competing attractor states (Litwin-Kumar and Doiron, 2012; Deco and Hugues, 2012; Lagzi and Rotter, 2015). Such assemblies and clusters increase the reliability of a potential readout by increasing spiking correlations, at the cost of reducing the dimensionality of the activity state space. The models cited above implement the mechanism simply by assigning a stronger connection weight or a higher connection probability to pairs of neurons belonging to the same cluster. However, biological neuronal networks have complex *higher-order* structure that goes beyond connection strengthening on a pairwise level, such as overexpression of triad motifs (Song et al., 2005b; Perin et al., 2011). Notably, this structure has been demonstrated to not be captured by models that only assign different strengths for different pathways, such as the ones cited above (Reimann et al., 2017a; Gal et al., 2017; Reimann et al., 2017b; Gal et al., 2019). Furthermore, Renart et al. (2007) pointed out the difficulty to obtain stable states at both a baseline and elevated firing rates that also have irregular spiking activity - as is observed in vivo - in large models with homogeneous connectivity. They pointed out that this may be rectified by more complex connectivity, specifically, one with hierarchical structure. This leads to the following question: Are there higher-order structures of connectivity in neuronal networks that support reliable network states, characterized by the activation of sets of assemblies, and transitions between them?

To find possible connectivity mechanisms for this purpose, we analyzed a biological connectome at cellular resolution (The MICrONS Consortium et al., 2021). Specifically, we detected and analyzed *directed simplices* (Reimann et al., 2017b) in the excitatory subnetwork. Directed simplices are large, tightly connected connectivity motifs with directionality giving them an input (source) and an output (target) side. They have been consistently demonstrated to be overexpressed in neural circuits of many organisms and at all scales, ranging from C. elegans (Reimann et al., 2017b; Sizemore et al., 2019; Shi et al., 2021) to rat cortical circuits (Perin et al., 2011; Song et al., 2005b; Reimann et al., 2017b), and to human regional functional connectivity (Sizemore et al., 2018). In simulation studies (Reimann et al., 2017b; Nolte et al., 2019) they have been demonstrated to affect network function by increasing spiking correlations and facilitating reliable information transmission from their input to output. Their abundance, strong internal connectivity and correlations make them relevant as a potential structural source of assembly formation, and the directionality of their connectivity a potential mechanism for the temporal transitions between them.

Inhibitory interactions may also play an import role. For example, competition between assemblies may be mediated by inhibitory interneurons (Bathellier et al., 2012). Inhibitory control has been shown to be crucial for a balanced, asynchronous state (Renart et al., 2010), but is often considered to be dense and non-specific (Fino and Yuste, 2011; Packer and Yuste, 2011; Litwin-Kumar and Doiron, 2012; Curto et al., 2019). In contrast, Schneider-Mizell et al. (2023) found types of specificity in inhibitory connectivity. This specificity was targeted at classes of neurons and their dendritic domains, independent of individual neuron identity. Znamenskiy et al. (2024) described specificity of inhibition at a more fine-grained level. They found that PV-positive neurons are more likely to connect to pyramidal cells that innervate them than expected. More generally, they proposed that PV-positive neurons connect predominately and more strongly to pyramidal cells with similar stimulus preference. In simulations, they demonstrated that this rule promotes a stable activity regime. In the context of assemblies, this rule implies inhibition that is targeted based on assembly membership, which has been found advantageous before in models (Rost et al., 2018). In that regard, Lagzi et al. (2021) predicted PV-positive neurons forming stabilizing inhibition within assemblies and SST-positive neurons competitive inhibition between assemblies, based on different forms of plasticity observed in the two classes. Due to the distributed nature of assemblies (Carrillo-Reid et al., 2015), such specificity is hard to measure without complete knowledge about which neurons participate in which assemblies. If directed simplices are correlates of assemblies, we may be able to find inhibitory specificity by analyzing connections between inhibitory neurons and these motifs, and specifically which neurons in these motifs are targeted.

At this level of complexity, the analyses have to be carefully controlled by comparing to relevant null models of connectivity, in order to understand the anatomical, morphological and molecular mechanisms giving rise to the results. If results can be explained by the spatial arrangement of neurons and distance-dependent connectivity, the case for their functional relevance is weak. At the same time, the shape of axons and dendrites has been shown to be a contributing factor to the complexity of connectivity (Stepanyants and Chklovskii, 2005) but it remains unclear to what degree these geometrical factors interact with other mechanisms affecting connectivity, such as activity dependent structural plasticity. Controls taking into account some mechanisms that shape connectivity but not others can help us disentangle the mechanisms.

As a data source of ground truth connectivity at the scale required for an exhaustive analysis, we used a connectome extracted from a freely available electron microscopic (EM) reconstruction of neural tissue, the MICrONS 1mm^3^ mouse visual cortex dataset (The MICrONS Consortium et al., 2021), from which we extracted locations of 60,048 neurons and their internal synaptic connectivity. We considered multiple centrally placed subvolumes to avoid edge effects and gauge the variability of the results. As controls we used random distance-dependent connectomes on the same neuron locations, and a recently released morphologically detailed model of cortical tissue with highly structured connectivity (Reimann et al. (2024); Isbister et al. (2023); from here on: *nbS1 model*) as a baseline taking into account anatomical and morphological factors, but not molecular or plasticity-related ones. That is, connectivity is derived from axo-dendritic appositions, hence taking the laminar placement of neurons and their neurite shapes into account. However, this does not recreate the preference of parvalbumin-positive interneurons to target dendritic locations close to the soma (Reimann et al., 2024), and represents a stochastic instance that is unaffected by rewiring through structural plasticity. Although it models a different organism (rat instead of mouse), age (juvenile instead of adult) and brain region (somatosensory instead of visual) than the MICrONS data, nbS1 served as a valuable control because it recreates the highly non-random overexpression of simplex motifs found in biological neural networks ranging from the worm to mouse (Egas Santander et al., 2024). This was crucial, as simplex motifs were at the core of all of our analyses. The approach of Udvary et al. (2022) could have served similarly well, as it derives connectivity from individual morphologies, which has been demonstrated to be required for non-random higher-order structure (Gal et al., 2019). To ascertain the functional relevance of directed simplices, we confirmed the *in silico* finding in Reimann et al. (2017b) that the activity correlation between connected neurons increases if the connection belongs to a simplex motif. To this end, we used calcium imaging data of neuronal activations to visual stimuli that was co-registered with the EM data. To determine the effect of potential reconstruction errors in the source data, we repeated analyses for a later version of the MICrONS data that was subjected to additional manual proofreading, and tested whether our results were stronger or weaker for the proofread neurons than the rest.

Conducting analyses as outlined above, we found: First, a hidden, lower-dimensional, directed and divergent network acting as a backbone of the recurrent connectivity and made up of excitatory simplices. Second, specific inhibition that is structured by the divergent network in that it follows the same direction. Third the non-random connectivity with the divergent network differed between inhbitory subtypes. Fourth, the inhibition in turn is under targeted control by specialized disinhibitory neurons. Fifth, these trends exist in the higher-order connectivity and are invisible on the pairwise level. Sixth, the trends are partially present in the computational model, indicating that they are morphologically prescribed and strengthened by other mechanisms. A plasticity-inspired rewiring rule makes the excitatory subnetwork of the model more similar to the EM connectome, but not the inhibitory to excitatory connectivity, indicating the presence of additional mechanisms. Seventh, errors in the source data do not explain the results, but rather weakened their strengths.

## RESULTS

### High-dimensional simplices reveal directed and divergent connectivity in the excitatory population

We analyzed the graph structure of synaptic connectivity of version 117 of the MICrONS dataset (The MICrONS Consortium et al., 2021). To assess its variability, we considered fifteen 500 *×* 300*µm* overlapping subnetworks (Figure 1A1). Note that the subnetworks comprised the neurons whose somata were within the defining rectangular prism and all synapses between them, even if those synapses were outside the prism. We compared to distance-dependent control models fit to the data and the connectivity of a subvolume of the *nbS1 model* (Reimann et al. (2024); Isbister et al. (2023), Fig. 1A2). Analyzed volumes of the nbS1 model were slightly smaller, at 300 *×* 300*µm*, to approximately match the neuron counts inside the volumes (14309 *±* 708 neurons for MICrONS vs. 13538 *±* 289 for nbS1).

**Figure 1.**
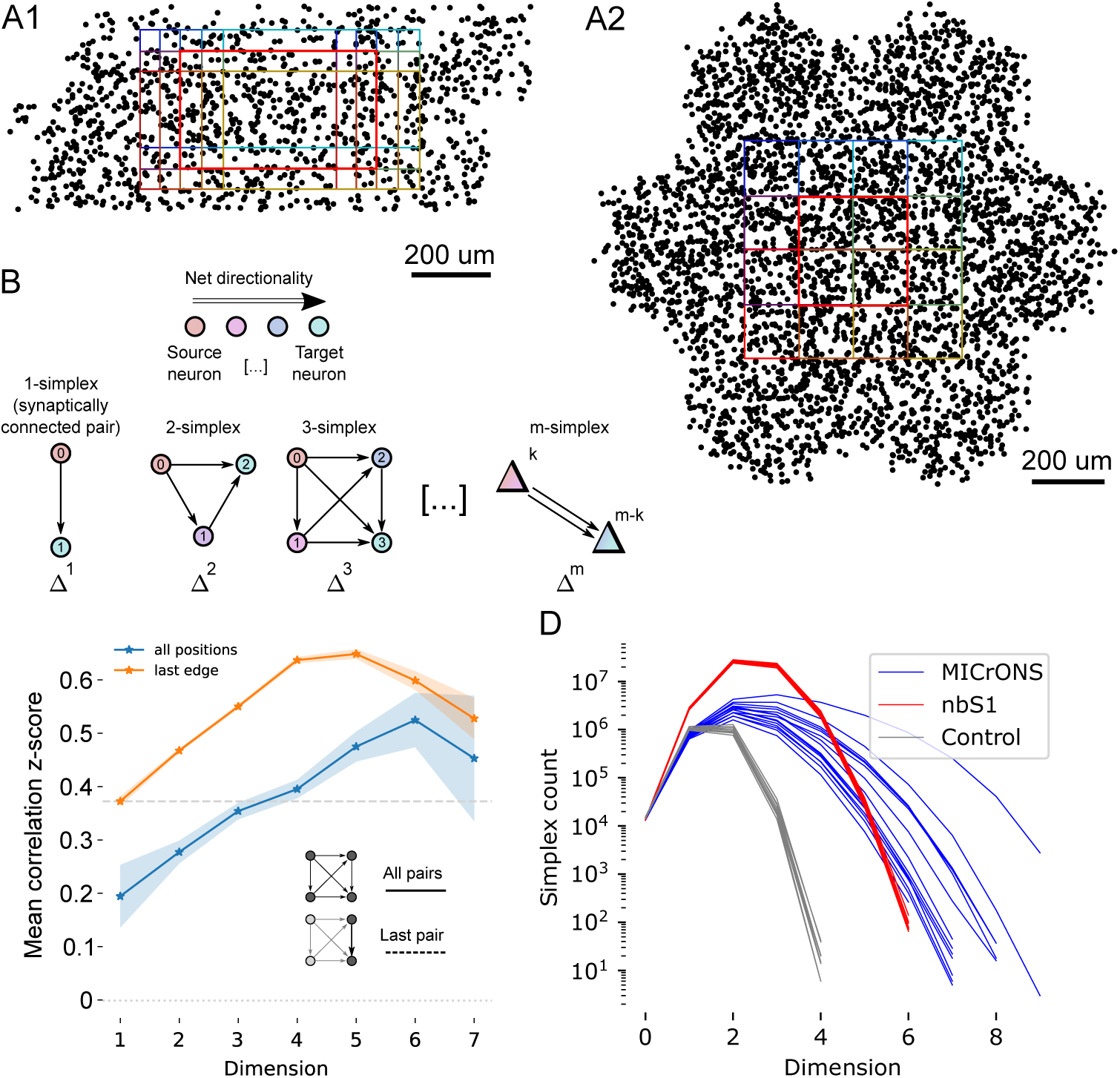
Analyzed connectomes and their simplicial structure. A1: Black dots indicate 2% of the neurons inside the *MICrONS volume*, an electron microscopic reconstruction of around 1*mm*^3^ of cortical tissue, the contained neurons and their connectivity (The MICrONS Consortium et al., 2021). Colored rectangles show the locations of 15 subvolumes separately analyzed. The thicker red outline indicates the *central subnetwork* that will serve as the example for some of the following analyses. A2: As A1, but for the *nbS1 volume*, a recently released, morphologically detailed model of cortical brain tissue (Isbister et al., 2023). Only 9 subvolumes were analyzed for the nbS1 volume. B: A *directed simplex* is a densely connected, feed-forward motif that generalizes the concept of an edge in a graph. An *m*-simplex consists of a *k*-simplex connected to an (*m−k*)-simplex, with all neurons in the first connected to those in the second, for any *m ≥* 0 and any 0 *≤ k ≤ m*. The *dimension* of a simplex is determined by the number of participating neurons. C: Mean z-scored correlations of the activity of pairs of neurons in the MICrONS data aggregated across repetitions against their simplex membership, shaded regions show the standard error of the mean. The x-axis indicates the maximum dimension over simplices the connection participates in. Blue: mean across all edges in simplices. Orange: values when only the last pair in a simplex is considered. Grey dashed line: Overall mean for connected pairs. Grey dotted line (lower): Overall mean for unconnected pairs. D: Simplex counts in the MICrONS and nbS1 volumes (blue and red) and control models with distance-dependent connectivity, fit to 9 of the MICrONS subnetworks (grey).

Analysis was conducted primarily with respect to *directed simplices* in the graphs representing the respective connectomes. A simplex with *dimension d* is a motif of *d* + 1 neurons with directed all-to-all connectivity (Fig. 1B). Each participating neuron has a *simplex position* numbered from 0 (the *source position*) to *d* (the *target position*). An edge must exist from neuron *a* to *b* if *b > a*; edges in the opposite direction can exist but are not required. As such, the concept generalizes the concept of a connection between neurons, which is a 1-dimensional simplex. In particular, all the nodes in a a simplex are distinct.

Simplex motifs have been demonstrated to be overexpressed in virtually all biological connectomes investigated (Perin et al., 2011; Song et al., 2005b; Sizemore et al., 2018). Moreover, they have been shown to have an impact on cortical activity, in particular, the spiking correlation of connected pairs of neurons increases with the maximal dimension of a the simplex the connection belongs to (Reimann et al., 2017b; Nolte et al., 2020). We confirmed both findings in the MICrONS data. Previous results on the same dataset found higher activity correlations for synaptically connected pairs of neurons (Ding et al., 2023). We extend this, showing even higher correlations when the connections are part of a simplex with a dimension above 4 or when it is placed at the target position of the simplex (Fig. 1C, S2). Simplex counts in dimensions above 4 were even higher than in the nbS1 model, but also more variable (Fig. 1D). This non-random structure resulted in overexpression of specific tirad motifs (Fig. S1). Both MICrONS and nbS1 had significantly increased simplex counts compared to the distance-dependent control. For an even simpler Erdos-Renyi control, i.e. a random assignment of connections to neuron pairs, independent of their distance, no simplices above dimension 3 emerged (not shown). We conclude that high-dimensional simplices have both structural and functional relevance in the MICrONS data and continue our analysis of them, with particular interest in their relationship to plasticity and inhibitory network specificity.

Individual neurons can participate in more than one simplex, forming complex, intersecting networks even when only simplices of a single dimension are considered (Fig. 2A). We calculated for the central subvolume of the MICrONS data the fraction of unique neurons separately for different dimensions and positions within the simplex (Fig. 2B, top). Results depended mostly on dimension considered, which is expected from the respective simplex counts being higher than the neuron count in some dimensions. When normalized with respect to the unique count for the source position, we found a strong increase of unique neuron count towards the target position (Fig. 2B, bottom). The strength of the trend mostly increased with dimension considered. This effect was observed consistently for all 15 subnetworks of the MICrONS volume (Fig. 2C1), and it was strongest in dimension 6. At the top dimension (7), results are noisy due to the relatively low motif counts. The trend was also found in the nbS1 model, but significantly weaker, peaking already at dimension 5 with a relative fraction of unique nodes of *<* 1.5 vs. *≈* 4.0 for microns (Fig. 2C2). It was extremely weak to non-existing in the distance-dependent control (Fig. 2C3).

**Figure 2.**
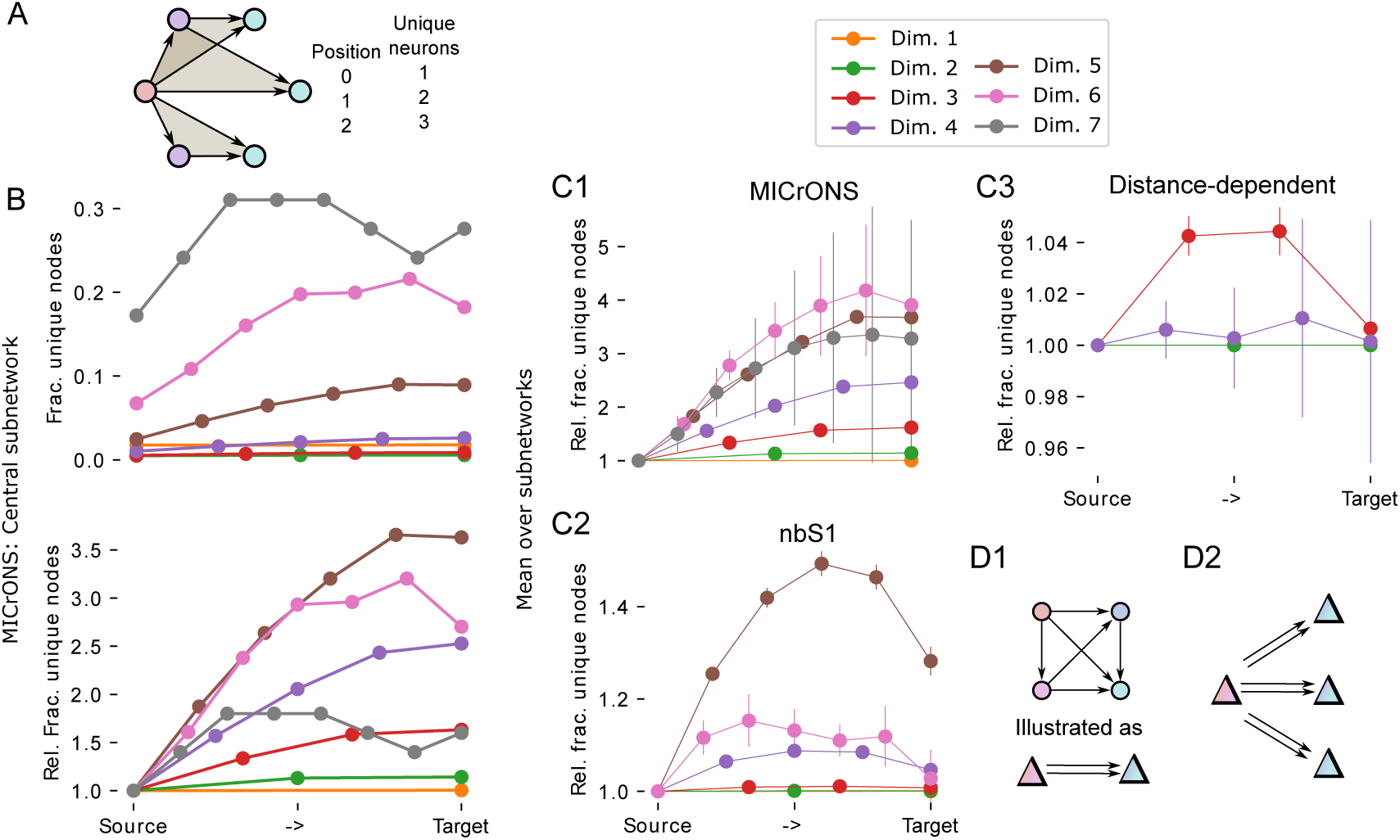
Analysis of simplices reveals divergence of directed information flow. A: As a neuron can be a member of several simplices, only a fraction of neurons in simplices of a given dimension is unique. The example shows three overlapping 2d-simplices (grey triangles). The table on the right lists the numbers of unique neurons in each position for the example. B: Top: Fraction of unique neurons in simplices of the central subnetwork of MICrONS, sorted by their position in the simplex from *source* to *target*. Bottom: Same, but normalized to fraction at the source. C1: As B, but means and standard deviation over the 15 subnetworks of the MICrONS data. C2,C3: As C1, but for the 9 subnetworks of the nbS1 model and distance-dependent control models. D: Schematic drawing interpreting the results. D1: As in Fig. 1B, we simplify the source and sink side of a simplex of any dimension (here 3-dimensional for illustration) to a red and a blue population, with a double arrow indication the direction of simplicial connectivity. D2: We found a higher-order connectivity motif with connectivity from a relatively small number of sources to a larger number of targets.

As connectivity in simplices is directed from the source towards the target neuron, this reveals aspects of the structure of information flow in the network: We found a highly non-random flow of connectivity from a compact source to a larger number of potential targets (Fig. 2D). Notably, this divergence is only visible when high-dimensional simplices are considered as it is entirely absent on the level of individual edges (Fig. 2, orange). We call the dimension where the divergent trend is strongest the *target dimension* of a network (6 for MICrONS, 5 for nbS1, 3 for distance-dependent; see Fig. 2C). Results for the following analyses will focus on and compare the respective target dimensions. Note that we do not intend to propose a special biological role for the target dimension, we only focus on it as a “target” for further analyses to keep the remainder of this work simpler to understand.

### Inhibition is structured by the simplicial structure of excitatory neurons

Next, we analyzed the graph locations of inhibitory neurons with respect to the simplicial structure (Fig. 3A). First, we found that in the central MICrONS volume excitatory neurons participating in 6d-simplices (i.e., the *6-core* of the central subnetwork) innervate and are innervated by more inhibitory neurons than the rest (Fig. 3B1). We confirmed that this was not merely an edge effect, where neurons in a more central spatial position are simply more likely to be connected to both excitatory and inhibitory neurons, as follows. First, while we detected simplices only for excitatory neuron inside the 500 *×* 300*µm* subvolume, we considered for this analysis inhibitory neurons of the entire MICrONS volume, thereby reducing the impact of the border drawn. Second, we compared the distances of neurons from the center of the volume to their inhibitory degrees, finding no dependence at all (Fig. S4). Third, neurons of the 6-core covered almost the entire range of distances from the volume center (Fig. S4, red vs. black). In the central nbS1 volume we conducted a corresponding analysis for its target dimension of 5, but since its 5-core contained more than half of the neurons, we also considered dimension 6 (Fig. 3B2). The increased inhibitory in-and out-degree was also present in the nbS1 model, but weaker: A 60% increase of inhibitory in-degree for neurons in the 6-core vs. 90% for MICrONS; a 100% increase of inhibitory out-degree vs. 200% for MICrONS.

**Figure 3.**
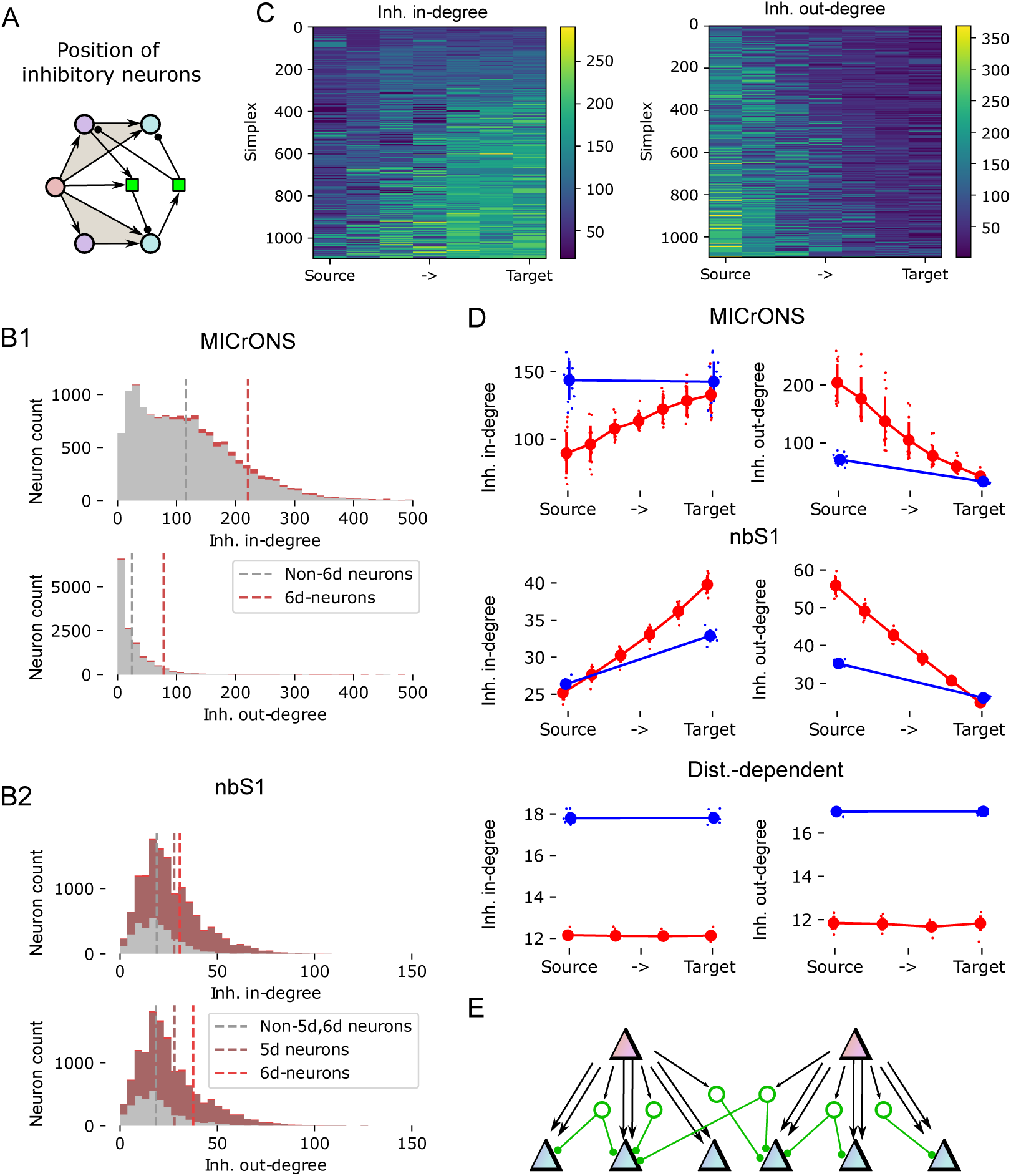
Non-random positioning of inhibition within the simplicial network. A: We consider the graph locations of inhibitory neurons (green) relative to simplices. Specifically, the number of inhibitory neurons innervating the neuron at a given position of a simplex (*inhibitory in-degree*) or innervated by it (*inhibitory out-degree*). B1: Distribution of the inhibitory in-and out-degrees of the central subnetwork of MICrONS. Red: For neurons participating in at least one 6d-simplex. Grey: For other neurons. Dashed lines: respective means. B2: As B1, but for the nbS1 model. Additionally, data for the 5-core is shown (dark red). C: Inhibitory in-degree (left) and out-degree (right) for neurons in all 1093 6d-simplices of the central subnetwork of MICrONS. Each row indicates one simplex, sorted by lowest (top) to highest (bottom) degree. Neurons participating in that simplex are indicated from left to right, sorted source to target. D: Mean inhibitory in-and out-degrees of neurons in each position of a simplex. Indicated are values for individual subnetworks (small dots) and mean and standard deviation over subnetworks (large dots and error bars). Red: data for the dimension with the highest directionality in Fig. 2C-E, i.e. 6 for MICrONS, 5 for nbS1, 3 for distance-dependent; blue: Data for 1d-simplices, i.e. source and target neurons of individual synaptic connections. E: Continuation of the schematic in Fig. 2D: Inhibition (green) is mainly excited by neurons in source positions of simplices (red) and innervates neurons in target positions (blue).

Crucially, this increase was strongly dependent on the position of a neuron in a 6-dimensional simplex (Fig. 3C). It was specifically neurons around the source position that innervated many inhibitory neurons, and neurons around the target position were strongly innervated by inhibitory neurons. Additionally, connections adhering to that principle were also formed by more synapses per connection (Fig. S3) indicating that such connections are selective strengthened through structural plasticity. Once again, the trend was consistently present in all 15 subnetworks of the MICrONS volume (Fig. 3D, top) and also present in the nbS1 data, but absent for the distance-dependent control (Fig. 3D, middle and bottom). Similar to the previous result of a divergent excitatory structure, the trend was almost undetectable when only single edges (1-simplices) are considered (Fig. 3D top, red vs. blue). Except for the nbS1 model, where it was visible but weaker when single edges are considered. We conclude that disynaptic inhibition between excitatory neurons is specific to the connectivity structures formed by high-dimensional simplices, and its directionality follows the direction given by the simplices (Fig. 3E).

### Inhibition implements specific competition between simplicial motifs, moderated by targeted disinhibition

Encouraged by the non-random targeting of disynaptic inhibition we found, we analyzed its structure further. We now considered the *structural strength of disynaptic inhibition* between pairs of simplices in the respective target dimensions. Specifically, we counted the number of *E→ I → E* paths where the first neuron was member of simplex *i*and the last neuron of simplex *j* (Fig. 4A). Note that this is a purely structural measure that does not take the physiological strengths of individual inhibitory synapses into account. The resulting matrix of structural disynaptic inhibition strengths between 6d simplices in the central subnetwork of MICrONS appeared highly structured and symmetrical (Fig. 4B1). To confirm this appearance, we compared it to a random control where we preserved the inhibitory in-and out-degree of each simplex, but shuffled their targets (Fig. 4B2). We found that the actual matrix contained values outside of the distribution given by the control, both significantly weaker and stronger (Fig. 4C, top). Repeating the analysis for all 15 MICrONS subnetworks we found strongly varying results (Fig. S5). Importantly, in each subnetwork we found data points both on the left and on the right side of the control. In some cases, this even manifested in two separate peaks, in others the peak on the right side of the control dominated. This demonstrates that disynaptic inhibition is structurally significantly weaker than expected between some pairs of simplices and significantly stronger between others, i.e., that it is targeted or specific. In terms of matrix symmetry, we computed the normalized variance of the values of the matrix minus its transpose (see Methods, Fig. 4D). The expected value of this measure for unstructured data is 2.0. Indeed, this was found in the shuffled control. The value for the actual data was significantly lower, indicating a high degree of symmetry. This finding was consistently present in subnetworks but varied in strength. Both specificity and symmetry were present for the nbS1 model in its target dimension 5, but weaker; and they were not present in the distance-dependent control (Fig. 4C, middle and bottom; 4D).

**Figure 4.**
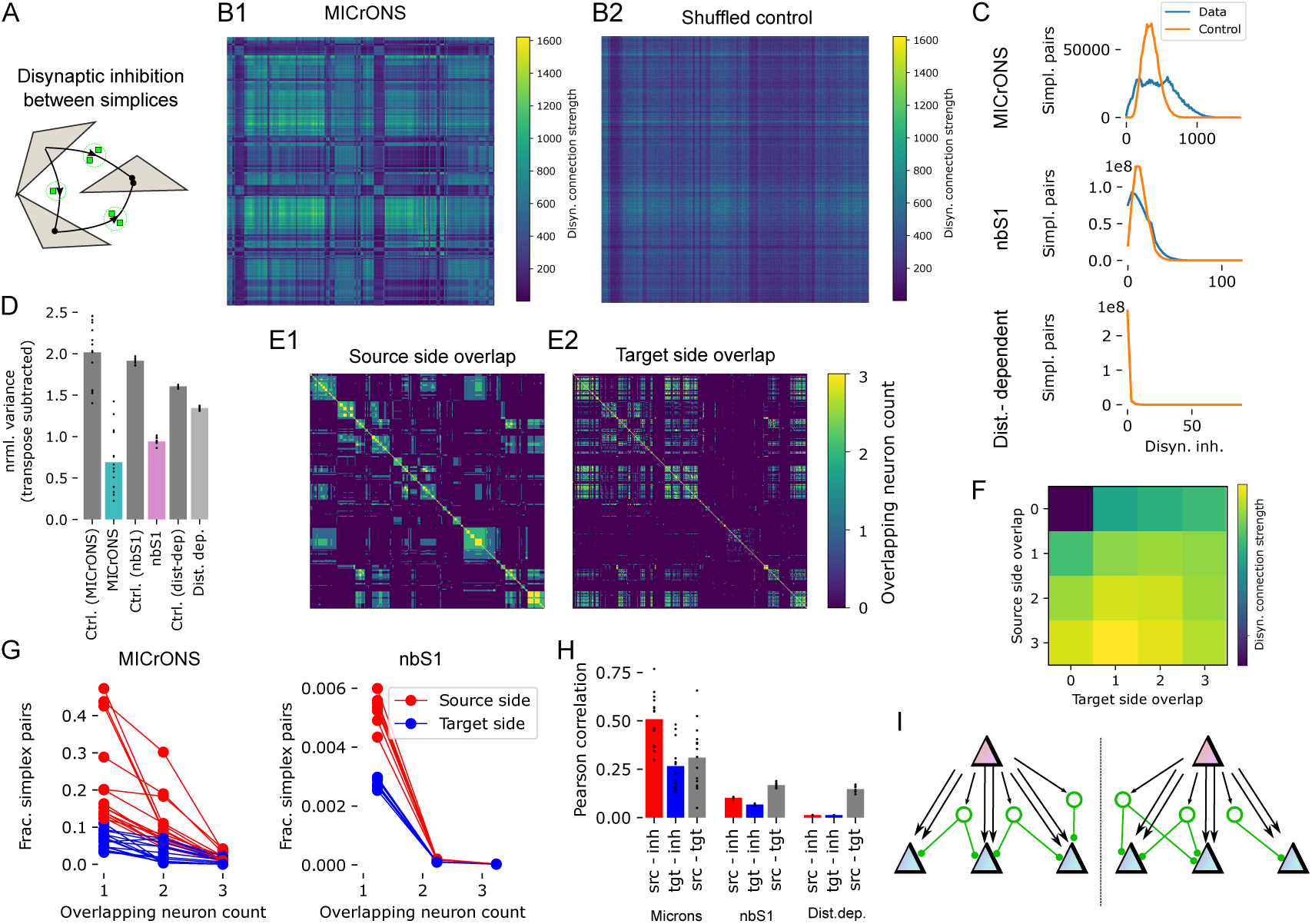
Specific, non-random disynaptic inhibition between simplices. A: We consider the strengths of *disynaptic inhibition* between pairs of high-dimensional simplices. B1: Disynaptic inhibition strength between pairs of 6d-simplices in the central subnetwork of MICrONS. Color indicates the number of paths from a neuron in the simplex along the vertical axis, to an inhibitory neuron, and then to a neuron in the simplex along the horizontal axis. B2: Same, for a random control, where the inhibitory in-and out-degrees of all simplices are preserved, but their sources and targets are shuffled. C, top: Distribution of disynaptic inhibition strength values in B1 (blue) and its control in B2 (orange). Middle: Same, for disynaptic inhibition between 5d-simplices in the nbS1 model. Bottom: For 3d-simplices in the distance-dependent controls. D: As a measure of symmetry of disynaptic inhibition matrices *M*, we show 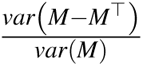. Dots indicate values for individual subnetworks, bars the means for (from left to right): Shuffled controls for MICrONS, MICrONS, shuffled control for nbS1, nbS1, shuffled control for the distance-dependent models, distance-dependent models. E: Number of neurons in common within the first three positions (*source side*, E1) and last three positions (*target side*, E2) of pairs of 6d-simplices in the central subnetwork of MICrONS. F: Mean disynaptic connection strength (as in B1) for pairs of simplices with the source-and target side overlap indicated along the vertical and horizontal axes. Mean over the 15 MICrONS subnetworks. G: Left: Normalized distribution of values in E1 (source side, red) and E2 (target side, blue). Right: Same for 5d simplices in the nbS1 model. Values missing from a sum of 1.0 are for pairs with an overlap of 0. H: Pearson correlations between source side overlap (as in E1), target side overlap (as in E2) and disynaptic inhibition (as in B1). Dots indicate values for individual subnetworks, bars the respective means. For (from left to right): MICrONS, nbS1 and the distance-dependent controls. I: Continuation of the schematic in Fig. 3E: The same simplices that excite inhibitory neurons from their source side (red) are also inhibited by them on their target side (blue).

This led to the question, what are the factors that affect disynaptic inhibition strength? What determined whether a value samples from the high or low peak? We considered *source side overlap* and *target side overlap*: The number of neurons a pair of 6d-simplices has in common in positions 0-2 (but in any order within that range; source side) and in positions 4-6 (target side; for details see Methods). This overlap tended to be stronger on the source side, as expected from the lower fraction of unique neurons in those positions (Fig. 4E, G). We found that the structural strength of disynaptic inhibition increased with both measures, but more so with source side overlap (Fig. 4F). This was confirmed by calculating the Pearson correlation between the three values (Fig. 4H). Results fluctuated over subnetworks but were consistently largest for correlation between source side overlap and disynaptic inhibition. Once again, the same trend was confirmed but weaker in nbS1 and not present in the distance-dependent control. These findings indicate that inhibition primarily implements a form of symmetric competition between multiple targets that are activated by the same source (Fig. 4I). Note that this differs from established models of competitive inhibition, where largely non-overlapping pools of excitatory neurons compete via inhibitory neurons. Since both source and targets are parts of the same simplices, this leads to a situation where the target neurons are simultaneously excited and disynaptically inhibited by the source population. This may facilitate a balance of excitation and inhibition, where minute fluctuations of inputs on top of a high-conductance state determine the outcome, which has been shown to lead to asynchronous activity as observed in vivo (Renart et al., 2010).

Another function of inhibitory connectivity is *disinhibition*, i.e., the inhibition of other inhibitory neurons. A subclass of Vasointenstinal protein (VIP)-expressing interneurons have been demonstrated to implement this function (Pi et al., 2013) and the presence of a class of inhibitory neurons specifically targeting other inhibitory neurons has been previously demonstrated in the MICrONS dataset (Schneider-Mizell et al., 2023). In the central subnetwork of MICrONS we found indication that the disinhibition preferentially targets the inhibitory neurons mediating competition between simplices (Fig. 5A, top): Not only were inhibitory neurons with many connections to and from 6-dimensional simplices more strongly inhibited themselves (Fig. 5A, middle), but this was specifically mediated by neurons specializing in such disinhibition. The fraction of inhibition coming from neurons with inhibitory targeting preference increased from 20% for neurons weakly interacting with simplices to 45% for the strongest interacting neurons (Fig. 5A, bottom right). We ensured that this is not an edge-effect by comparing the inhibitory indegrees to the distance from the center of the analyzed volume, finding no significant correlation (Fig. S6).

**Figure 5.**
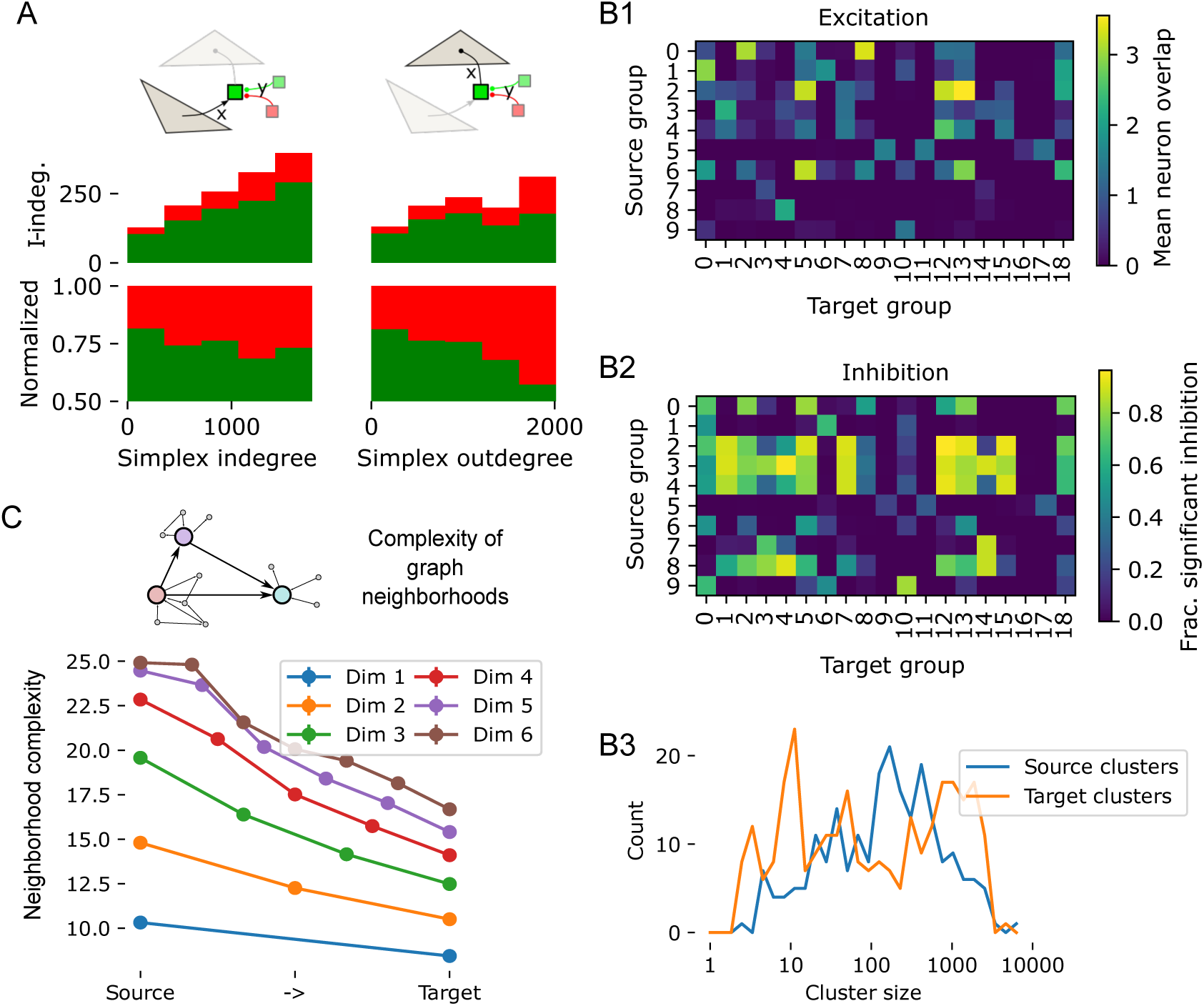
Specific disinhibition and lower-dimensional structure of the competitive simplicial network. A, top: We analyze for neurons participating in lateral inhibition between simplices (opaque green square) how they are themselves inhibited (transparent squares). We consider for the x-axis the *simplex degrees* of inhibitory neurons in the microns data. That is, its number of connections from (left, simplex indegree) or to (right, simplex outdegree) 6-dimensional simplices. We consider for the y-axis their indegree from inhibitory neurons with significant targeting preference for other inhibitory neurons (*p ≤* 10*^−^*^6^; 15% of inhibitory neurons; red), or from the remaining inhibitory population (green). Stacked bar plot for absolute (middle) or normalized (bottom) inhibitory indegrees. B: Strengths of excitation (B1) and inhibition (B2) between groups of 6d-simplices in the central subnetwork of MICrONS. Groups were derived by clustering the source side overlap matrix (see Fig. 4 D, left), yielding *source groups* along the vertical axis, and by clustering the target side overlap matrix (Fig. 4D, right), yielding the *target groups* along the horizontal axis. For excitation, we considered the mean number of overlapping neurons in any position over pairs of simplices in the indicated groups. For inhibition we considered the fraction of pairs with disynaptic inhibition strength higher than the 95th percentile of the corresponding shuffled control (Fig. 4A, B). B3: Distributions of source and target group sizes in the 15 MICrONS subnetworks. C: Complexity of subgraphs given by graph neighborhoods of individual nodes in the central subnetwork of MICrONS. Indicated are the mean values for neighborhoods of nodes participating in the indicated position of simplices of the indicated dimension. For details on the complexity measure used see Egas Santander et al. (2024), Methods.

### Spatial and topological structure of disynaptic inhibitory competition

So far, we have found a higher-order network acting as a backbone of the recurrent connectivity with divergent feed-forward excitation from groups of source simplices to groups of target simplices and specific lateral inhibition between them. Here, we describe the structure of that feed-forward network and the neurons participating in it.

As we have demonstrated that overlap in source and target neurons shapes the structure of disynaptic inhibition between simplices, we began by detecting these overlapping groups that inhibit or are inhibited together. We determined *source* and *target groups* of simplices by clustering the matrices of source side and target side overlap respectively. We used the Louvain algorithm, which aims to maximize *modularity* of the resulting clusters, i.e. the strengths of connections (in our case: overlap) within a cluster compared to connections across clusters. Next, we estimate the strength of excitation and inhibition between source and target groups based on all pairs of simplices *s*_1_*, s*_2_ where *s*_1_ is member of the source group and *s*_2_ of the target group. The aim was to characterize to what degree that connectivity of the source-and target group was characterized by the trends we have found so far. For excitation, this was participation in overlapping simplices (Fig. 2). Hence, we calculated the mean overlap of neurons in any simplex location (Fig. 5B1). For inhibition, this was providing disynaptic inhibition between simplices that is stronger than expected by chance (Figs. 3, 4). Hence, we calculated the fraction of pairs with significantly strong inhibition, i.e., with a disynaptic inhibition strength above the 95th percentile of the corresponding shuffled control (Fig. 5B2).

The result was a 10 *×* 19 connectivity matrix of a feedforward network with relatively sparse excitation and specific inhibition. We note that the numerical values of our measures of excitation and inhibition are not directly comparable, but conclusions may be drawn based on their relative values. Aggregated over all 15 MICrONS subnetworks, the number of source and target groups varies, as does the number of simplices belonging to them. Indeed, the number of simplices in a cluster falls into a wide distribution spanning almost four orders of magnitude, indicating an organization principle at multiple scales (Fig. 5B3).

Taken together, this raises the question: what is the role of the complex higher-order structure within the feedforward structure? If feedforward connectivity is called for, why not implement a simple feedforward network? Simulation results (Egas Santander et al., 2024) predicted a role of higher-order structure in increasing the reliability of the network response. Egas Santander et al. (2024) characterized the graph neighborhoods of neurons in terms of how much their connectivity deviates from random, for example, with overexpression of bidirectional connectivity. They found that the presence non-random neighborhoods increases reliability globally, while more random neighborhoods provide better information readouts. Considering the feedforward structure given by high-dimensional simplices in our results, we expect the following: As target neurons of simplices are the outputs of the circuit, we would expect them to be members of neighborhoods providing good information readout. Conversely, on the input side given by source neurons, we would expect subnetworks facilitating a reliable response. Indeed, replicating the analysis of Egas Santander et al. (2024) (Fig. 5C), we found the neighborhoods of target neurons to be more similar to random graphs than those of source neurons, with the strength of this trend increasing with simplex dimension. Taken together this indicates that indicates that the purpose of the complex simplicial structure is to channel activity from parts of the network at the input side of the motif and that improve spiking reliability into other parts that facilitate efficient readout.

By considering all neurons participating in at least one simplex of a source group, we can visualize its internal connectivity (Fig. 6A). This illustrates the dense feed-forward connectivity given by the 6-dimensional simplices contained in the group, but also the more complex, highly recurrent nature of this network: 38% of edges between nodes do not participate in the simplices making up this source group. Over all source groups, this is 41% *±* 10% (mean *±* std). This highlights once more how the trends described in this manuscript exist on the higher-order level of structured connectivity between neuron motifs and not individual pairs. So far, we have soundly refuted the idea that any of the observed trends are caused merely by the relative locations of neurons using distance-dependent control connectomes. However, we still wanted to see to what degree neurons in a source group cluster together spatially. We found neurons in a source group span up to 500*µm* (Fig. 6B, S7). Furthermore, groups can overlap spatially despite only marginal overlap in participating neurons (e.g. groups 0 vs 2 in Fig. S7). Information flow in local circuits such as these has been traditionally described as a flow between cortical layers (Felleman and Van Essen, 1991). Conversely, we found no layer specificity, with neurons in source positions (Fig. 6B, blue) and target positions (Fig. 6B, red) being present in all layers. This highlights once more that our results reveal a higher-order structure that is largely invisible at the pairwise level. Curiously though, we found that pairs of source neurons tended to be further apart from each other than target neurons (average of 220*µm* for pairs of source vs. 130*µm* for target neurons; Fig. S8). This may be explained by the source position containing more neurons classified as 5P IT (Table 1), a class that is known to form long-range corticocortical connections (Harris et al., 2019). Another type overexpressed in the source position is 4P (layer 4 pyramidal cells), which is consistent with the classical view that they are the inputs of a local circuit (Felleman and Van Essen, 1991). Conversely, the 6IT class formed the outputs, participating almost exclusively in the target position.

**Figure 6.**
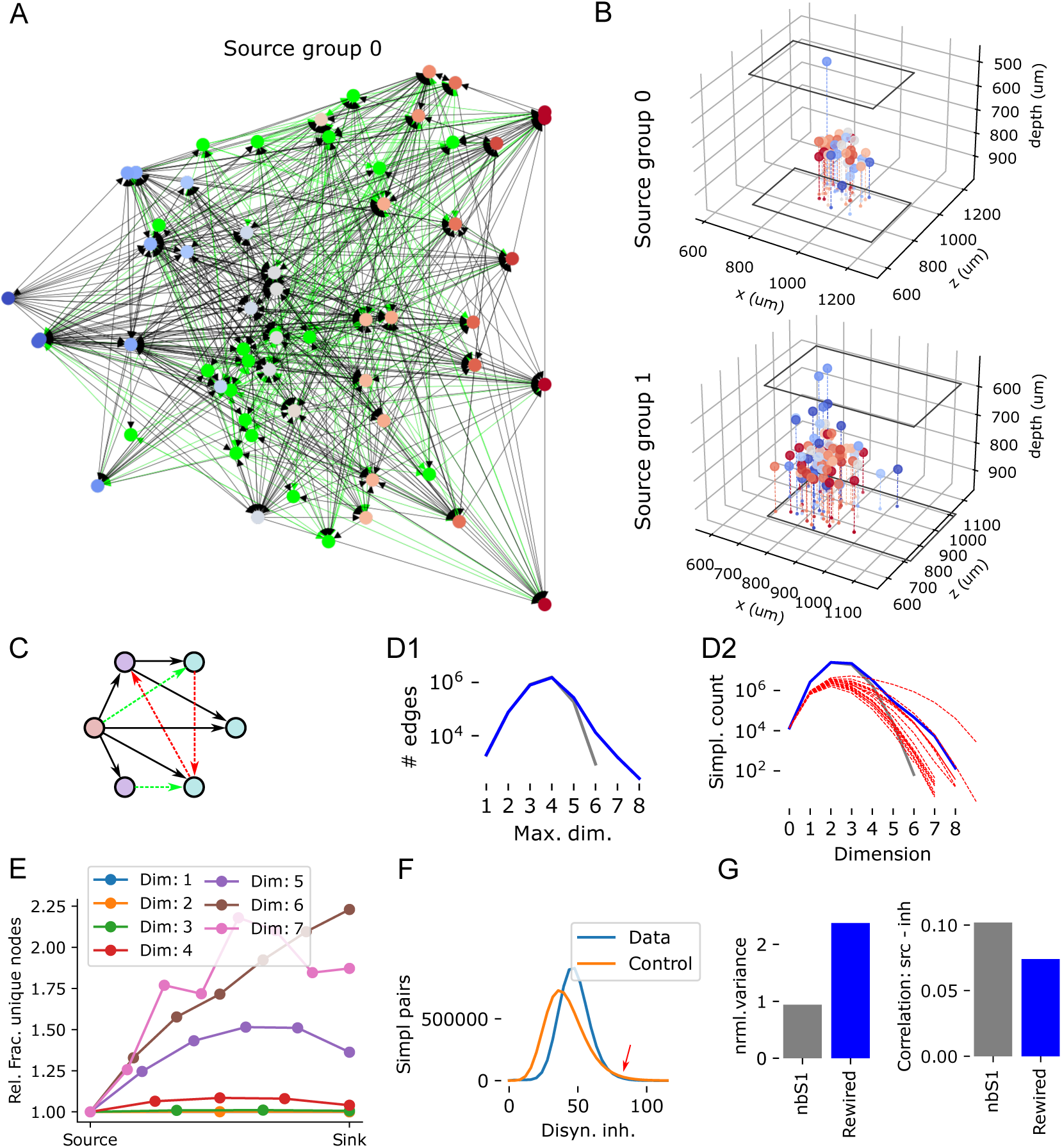
A rewiring rule to better capture the non-random structure of connectivity. A: Example connectivity of a source group (group 0 in Fig. 5B) Excitatory neurons that are member of any simplex in the group are placed along the horizontal axis and colored from blue to red, based on their mean position in a simplex. In green: 20 randomly picked inhibitory neurons connected to the simplex group. B: Spatial locations of neurons in source groups 0 and 1. Black outline indicates boundaries of the central subnetwork of MICrONS. See also Fig. S7). C: A rewiring rule removes edges participating only in low-dimensional simplices (red) and adds edges that are likely to form new high-dimensional simplices (green). D1: Maximum dimension of simplices an edge is participating in. Grey: nbS1 model; blue: nbS1 model with 1.2% of edges rewired. D2: Simplex counts as in Fig. 1D. Red dashed lines indicate results for the various MICrONS subnetworks. E: Divergence in terms of the number of unique neurons as in Fig. 2C for the rewired nbS1 model. F: Histogram of disynaptic inhibition strengths as in Fig. 4C for the rewired nbS1 model. Red arrow indicates a region on the right side of the distribution where the value for the control is above the data. G: Symmetry of disynaptic inhibition (left; as in Fig. 4D) and correlation between source side overlap and disynaptic inhibition strength (right; as in Fig. 4H), comparing the nbS1 model (grey) to its rewired version (blue).

**Table 1.** 6-dimensional simplex composition in terms of cell types. Below divider: For comparison, the fractions of neurons of each cell type, and the fractions of outgoing and incoming edges associated with each cell type.

|  |  | Class |  |  |  |  |  |
| --- | --- | --- | --- | --- | --- | --- | --- |
|  |  | 23P | 4P | 5P_IT | 5P_PT | 6CT | 6IT |
| Position | 0 | 0.91% | 31.5% | 25.7% | 33.8% | 7.7% | 0.3% |
|  | 1 | 9.2% | 30.6% | 26.1% | 30.9% | 0.2% | 3.0% |
|  | 2 | 12.2% | 33.6% | 18.5% | 30.6% | 0.0% | 5.0% |
|  | 3 | 11.9% | 30.2% | 19.6% | 30.3% | 0.1% | 7.9% |
|  | 4 | 8.9% | 29.6% | 8.5% | 35.2% | 0.1% | 17.7% |
|  | 5 | 8.8% | 23.2% | 8.5% | 44.3% | 0.0% | 15.2% |
|  | 6 | 7.2% | 23.0% | 7.1% | 39.1% | 0.0% | 23.0% |
| Neurons |  | 28.7% | 29.3% | 12.3% | 3.5% | 16.6% | 9.5% |
| Out-edges |  | 27.1% | 33.2% | 16.2% | 5.7% | 10.3% | 7.5% |
| In-edges |  | 34.4% | 31.1% | 13.4% | 6.1% | 9.0% | 6.0% |

### Potential rewiring mechanisms shaping connectivity

While present in the nbS1 model, the non-random connectivity features were consistently weaker than in the MICrONS data. One of the mechanisms missed by the model explaining the difference could be structural plasticity. Recent simulation results predicted that plasticity favors connections that participate in many high-dimensional simplices, and this has been confirmed in the MICrONS data (Ecker et al., 2023). Accordingly, we implemented a simple heuristic algorithm for rewiring excitatory connections that removes edges that only participate in low-dimensional simplices, and places edges that are likely to form new high-dimensional simplices (Fig. 6C). We chose to study this particular rule over others that have been proposed because it depends on the participation in simplices, which are at the core of our results. To give rise to the observed increase in unique neurons towards the target (Fig. 2C), we focused on placing edges between neurons on the source side (for details, see Methods). We found that rewiring of only 1.2% of connections succeeded in increasing edge participation and simplex counts significantly (Fig. 6D) with a divergent structure at higher dimensions (Fig. 6E). However, this excitatory rewiring alone reduced the specificity of disynaptic inhibition (Fig. 6F, G) between the simplices.

### Validity of the results

While connectomics based on electron microscopy can be extraordinarily accurate, it relies on automatic reconstruction algorithms that are imperfect. To yield correct connectivity for a neuron its dendrite and axon need to be traced throughout the entire volume. This process can be subject to false splits and false mergers. Additionally, synapses in the volume must be assigned to the correct dendrites and axons. This process is more error-prone for inhibitory or shaft synapses, with lower reported values for precision and recall (between 78% and 97% of values for excitatory or spine synapses; Motta et al. (2019); Shapson-Coe et al. (2021)). Such errors could affect our results, leading to the question of the validity of our findings. First, we would like to point out that our results indicate a highly non-random structure of connectivity that is extremely unlikely to emerge from random reconstruction errors. For synapse assignment in the dataset we used, precision of 96%, recall of 89% with partner assignment accuracy of 98% compared to manual identification was reported (The MICrONS Consortium et al., 2021). Degradation of the connectivity data within these bounds is mathematically incapable to provide an explanation for the non-random features we found. For example, to explain the presence of seven-dimensional simplices we report in Fig. 1D by chance, a graph without higher-order structure would have to be subjected to a degradation of at least 94% of connections (corresponding to only 6% recall, see Methods). And even in that case, the degradation, i.e. the set of missed synapses, would have to be selected with incredible specificity, as the probability that a 7d simplex survives the degradation by chance is 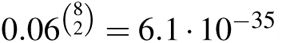. Similar numbers are found when one tries to explain the structure through false positives.

To further confirm the validity of our findings and understand the impact of reconstruction errors on our results, we tended to a later version of the MICrONS data (v1181 vs. v117 for the results above), that was subjected to manual proofreading of around 2% of the neurons (Fig. 7A. First, we repeated the previous analyses on the new data. We found that simplex overexpression was at the same level (Fig. Finally, we used variants of the analyses that we could run separately for neurons based on their proofreading status (see Methods). We found that neuron that passed proofreading participated in a larger number of simplices than neurons that were not proofread, or neurons where the axon failed proofreading (Fig. 7B, top). This was in part due to their higher degree, but also persisted after normalization for that effect (Fig. 7B, bottom). Or findings in Fig. 2 show that a smaller number of unique neurons participate in the source position of large simplices than in the sink position. While proofread neurons were overall more likely to participate in 6d-simplices than non-proofread ones, the fraction still increased from source to sink position (Fig. 7C, top). After normalization, the trend was roughly equally strong for proofread and non-proofread neurons, but absent for neurons that failed proofreading (Fig. 7C, bottom). Our results in Fig. 3 show that inhibitory neurons were more likely to be innervated by source neurons and more likely to innervate sink neurons. We found that proofread inhibitory neurons were overall more likely to interact with simplices in either direction and any position than non-proofread ones (Fig. 7D, E, top). The relative increase towards the sink position in one direction and the relative decrease in the other direction was stronger for proofread neurons (Fig. 7D, E, bottom). In summary, in all cases the non-random trend we found was stronger or as strong when only neurons that passed proofreading were considered. We conclude that our results presented in Figs. 1-3 are likely to underestimate the true strength of the reported connectivity trends. Therefore, we believe that the results in Figs. 4-5, which further explore and describe the proposed non-random structure, do not require additional validation.

**Figure 7.**
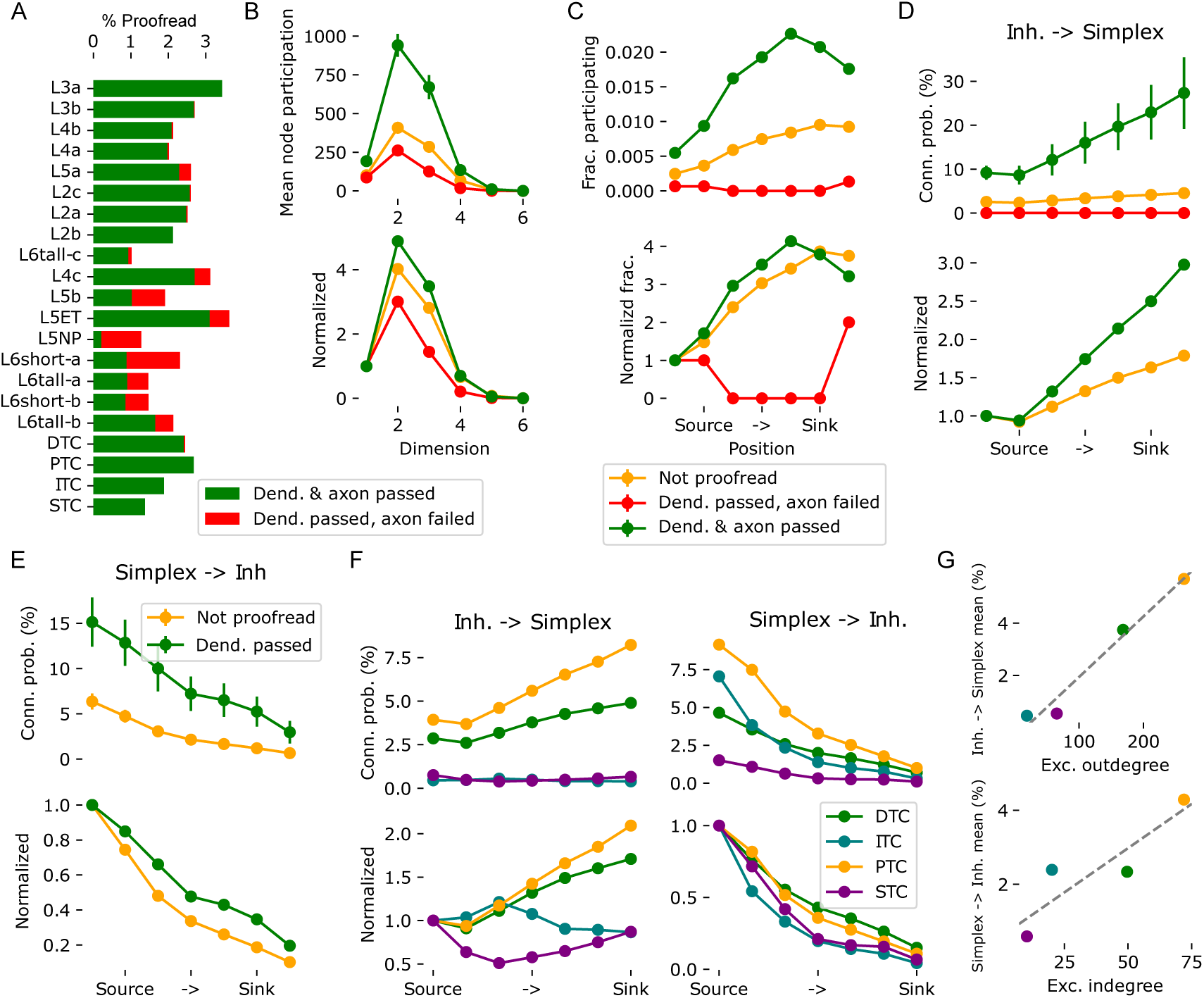
Validation and roles of inhibitory subtypes analyzed using a later revision of the data. A: Percentages of neurons proofread in version 1181 of the MICrONS data. Missing percentages were not proofread. The combination of dendrite failed and axon passed did not occur. B: Top: Mean number of simplices an excitatory neuron participated in (node participation) in various dimensions based on their proofreading status. Error bars indicate standard error of mean. Bottom: Same, but normalized against participation in 1d-simplices, i.e. neuron degree and without error bars. C: Fraction of excitatory neurons participating at least one 6d-simplex in the indicated positions, grouped by their proofreading status. Bottom: Normalized by the fraction for the source position. D: Probability that an inhibitory neuron innervates a 6d-simplex neuron in the indicated position, grouped by the proofreading status of the inhibitory neuron’s axon. The plot indicates the mean over simplices, error bars the standard error of mean. Bottom: Normalized by the value for the source position. Failed axons not indicates as the normalization led to division by zero. E: Probability that a 6d-simplex neuron in the indicated position innervates an inhibitory neuron, grouped by the proofreading status of the inhibitory neuron’s dendrite. Mean over simplices and standard error of mean indicated. Note that no dendrites failed proofreading. Bottom: Normalized as in D. F: Left / right as D and E, but instead of the proofreading status the cell type of the inhibitory neurons is considered. G: For inhibitory cell types the mean outdegree (top) and indegree (bottom) to / from excitatory neurons is plotted against the respective means over simplex positions in F. S9A), divergence reached the same level, althought the roles of dimension 5 and 6 were flipped (Fig. S9A). Simplicial targeting by inhibition (Fig. S9C remained unchanged. Its symmetry increased and its targeting was even better explained by source group overlap (Fig. S9D, E).

### Roles of inhibitory subtypes

The later version of the MICrONS data used for validation in the previous section also contains annotations of inhibitory neuron classes according to the schema proposed by Schneider-Mizell et al. (2023). The subtypes were labelled throughout the volume using a soma and nucleus feature trained metamodel (Elabbady et al., 2022). This allowed us to test whether these subclasses also interact with large simplices differently. Briefly, Schneider-Mizell et al. (2023) grouped inhibitory neurons into four connectivity classes: DTC or distally targeting cells are tentative SST-positive neurons, such as Martinotti Cells; ITC or inhibitory targeting cells are tentative VIP-positive neurons; PTC or proximally targeting cells are tentative PV-positive neurons, such as Basket Cells; STC or sparsely targeting neurons are tentative Neurogliaform Cells. We found that only DTC and PTC neurons innervated simplices with significant probabilities that increase from source to sink (Fig. 7F, left). This is not a surprise, as ITC neurons are defined by their low likelihood of innervating excitatory neurons and we only considered simplices of excitatory neurons. STC neurons are tentative Neuogliaform Cells, which have been associated with volumetric, i.e. untargeted, synaptic transmission. PTC cells innervate simplices slightly more strongly than DTC cells, with a higher preference for sink neurons, but the statistical significance of this distinction remains unclear (see below). Simplices innervated the subtypes at different levels (Fig. 7F, right) with a preference for source neurons present in all classes, but slightly stronger for STC and ITC neurons.

In general, the likelihood that a neuron innervated or was innervated by a 6d-simplex neuron in any position could be well predicted from the mean excitatory out-or in-degrees of the four classes (Fig. 7G). One exception was that ITC neurons were more likely to be innervated by simplices than DTC neurons despite having less than half their mean excitatory in-degree. This could indicate that source neurons of simplices are also specifically activating disinhibition by targeting a class of neurons that inhibits other inhibitory cells. However, we would like to point out that we were unable to assess the statistical significance of any of the results presented in Fig. 7F and G. This was because the inhibitory subclasses were proofread at different levels (Fig. 7A), which could explain the differences observed. Unfortunately, the number of proofread neurons was too low to make a double distinction between proofreading status and inhibitory subclass simultaneously. Yet, we would like to point out that the apparent stronger innervation of ITC neurons than DTC neurons by simplices cannot be explained this way, as ITC neurons were proofread at a lower rate than DTC neurons.

Finally, we wanted to be able to interpret our results in the context of a recent finding that PV-positive interneurons connect specifically to pyramidal cells that innervate them (Znamenskiy et al., 2024). As such, we investigated connection probabilities between interneuron -L2/3 PC pairs that were proofread and within 50*µm* of each other. This analysis was conducted on the entire volume. For the PTC class of neurons, that are thought to correspond to PV-positive cells (Schneider-Mizell et al., 2023), the degree of overexpression of bidirectional connectivity matched the reference (Fig. S10). Note, however, that the PC to PV-positive connection probability was lower (47% vs. 73% reported by Znamenskiy et al. (2024)). The overexpression of bidirectionality was lower for the DTC class and absent in ITC and STC.

## DISCUSSION

We analyzed the connectivity of an electron microscopic reconstruction of a 0.65mm^3^ volume of mouse visual cortex. This is considered the gold standard in cortical connectomics due to the size of the EM-reconstructed volume encompassing whole dendritic morphologies and allowing the analysis of local connectivity. Using methods of algebraic topology suited for discovering higher order connectivity motifs, we found an underlying feed-forward network with specific lateral inhibition implementing a mechanism of competition between different outputs. In turn, we found the lateral inhibition is under the control of targeted disinhibition from a specialized class of inhibitory neurons. No aspect of the structure was evident when first-order connectivity between individual neuron pairs was considered; it required specific techniques from the field of algebraic topology (in this context: *neurotopology*) that consider connectivity in motifs of any size, i.e., beyond purely pairwise. This contrasts this work with previous analyses of the same dataset. Schneider-Mizell et al. (2023) found important pairwise principles of inhibitory connectivity at subcellular resolution. Ding et al. (2023) quantified how aspects of neuron function, such as correlations relate to pairwise connectivity. Here, we have demonstrated that increases of correlations are stronger when participation in directed simplices, i.e., densely connected feedforward motifs, is considered. To allow other researchers to conduct these analyses efficiently, we have open-sourced the code in the form of the conntility and connalysis python packages (Egas Santander et al. (2024); see data and code availability statement).

While impressive, electron-microscopic connectomes are imperfect. Yet, they have a proven track record, enabling important discoveries relating to both excitatory and inhibitory connectivity (Motta et al., 2019; Shapson-Coe et al., 2021; Schneider-Mizell et al., 2023; Ding et al., 2023). We have demonstrated that synapse assignment errors at the rates reported for the dataset we used are mathematically impossible to explain our results. To account for another source of potential errors, false splits and mergers during tracing, we conducted specific validation analyses. We were able to demonstrate that these errors cannot explain our findings and the strength of our results would likely be even stronger in their absence. Our analyses may appear complex, and complex analyses are often brittle against data imperfections. But as they take an entire connectome with around a million edges into account, the results hold at the highest levels of significance. This is further demonstrated by the fact that the results were apparent even in the version of the source data that was subjected to very little proofreading. Another possible confounding factor of the type of analysis we employed is the presence of an edge effect, i.e. neurons in the periphery of an analyzed volume missing connections from outside the volume. In that regard, we have minimized its impact by considering connections from outside an analyzed subvolume, and by carefully evaluating and rejecting the null hypothesis that an edge effect explains a result. But most importantly, an edge effect would be equally visible in the pairwise connectivity, where we have demonstrated its absence. Thus, the presence of this structure is strongly supported by the data, even given its flaws.

While the electron-microscopic volume we considered was the largest to date, it still only spanned around 600*µm* in one direction. This means that we only considered a fraction of the inputs and outputs of the neurons we analyzed. For excitatory neurons, the maximum distance was further reduced by selecting subvolumes. Stepanyants et al. (2009) estimated the severity of the loss of connectivity to slicing. For excitatory connections in a 300*µm* slice, similar to our subnetworks, 90% of connectivity was found to be missing. Note that this estimate is for cat and the value for mouse or rat is likely lower due to the overall smaller scale of the brains. For inhibitory connections, the loss is much lower. We always considered all inhibitory neurons, even the ones outside the subvolumes to avoid edge effects. For 600*µm* slices corresponding to that, less than half of the inhibitory connectivity would be missing according to Stepanyants et al. (2009). It is difficult to say what the loss means for the simplicial structure of connectivity. Reimann et al. (2024) predict that inclusion of long-range connectivity increases the maximum simplicial dimension to 18 with a relatively small number of unique neurons in layer 4 and 5 acting as sources and a larger number of neurons, mostly in layer 5 acting as sinks. This prediction, which is based on the nbS1 model we used as a control in this work, is in line with our main finding of divergence of the excitatory network.

As a compromise between studying networks that are large enough to contain high-dimensional simplices, but still being able to assess the variability of results, we analyzed subnetwork that overlapped in their neurons. Their results were therefore not strictly independent. On the other hand, to affect the results, all nodes of a simplex had to be contained in a subnetwork. The overlap in simplices was therefore smaller than the overlap in neurons. Regardless, the overlap means that caution must be taken when interpreting our results quantitatively. The presence of the described structures is unaffected though, as they were consistently present in every subnetwork.

The non-random features of connectivity we found are possibly related to neural manifolds (Gallego et al., 2017; Stringer et al., 2019), the concept that circuit activity states exists and move around in a comparatively low-dimensional state. In this case, the state would be given by the combination of structurally determined source and target groups that are active, and the flow of information between them. Source and target groups also align with the concept of assemblies (Hebb, 1949; Harris et al., 2003; Dragoi and Buzsáki, 2006; Lopes-dos Santos et al., 2013), i.e., groups of neurons with correlated spiking activity. As neurons participating in a target group are excited as members of the same set of densely connected feed-forward motifs, they are likely to fire together. The principles of organization described are mostly orthogonal to the concept of information flow between layers, as all layers participate in all simplex positions. Still, our results were in line with established neuron roles: Both 4P and 5P IT neurons prefer to act in a source position (Table S1). This is in line with their established roles as initial inputs to local (4P) and distal (5P IT) cortical circuits (Felleman and Van Essen, 1991; Harris et al., 2019). Our results reinforce the idea of layer 6 as an output layer.

Within this structure, inhibition implements competition between different potential outputs. Such inhibition has been found to be crucial to achieve a balanced, asynchronous state (Renart et al., 2010), but is often considered to be non-specific blanket inhibition (Fino and Yuste, 2011; Packer and Yuste, 2011). In contrast, Rost et al. (2018) demonstrated that structure in inhibitory connectivity facilitates switching between assemblies without saturating their firing rates. Our results are more in line with the latter result, but additionally describe the internal structure of excitatory assemblies and specific positioning of inhibition within them. Additionally, we found indication, albeit non-conclusive, that disinhibitory (ITC, inhibitory targeting interneurons) neurons were targeted with even higher specificity by source neurons of simplices than other inhibitory types. This would mean that the lateral inhibition is activated at the same time as the neurons that serve to deactivate it, leading to a balance of excitation and inhibition also for inhibitory neurons. All of this together indicates an inhibitory network where the exact identity of source and target neurons matters, and that plays a crucial role in the computations implemented by the local circuitry. For a different class of neurons (PTC, proximally targeting cells, tentatively PV-positive cells) we confirmed a recent finding that they form more bidirectional connections with pyramidal cells in layer 2/3 than expected (Znamenskiy et al., 2024). This shows that their results are compatible with ours, as they coexist in the same network. More generally, Znamenskiy et al. (2024) proposed that PV-positive neurons innervate pyramidal cells that have the same stimulus response properties as them. This is compatible with the characteristic motif we found of a source neuron innervating an inhibitory neuron, which innervates the sink neuron. As they are tightly connected, the source and sink neuron of a simplex are likely to have similar response properties. In addition to this role of PV-positive neurons, Lagzi et al. (2021) predicted SST-positive neurons to mediate competitive inhibition between assemblies. In contrast, we found no difference in how PTC and DTC neurons (corresponding to PV-and SST-positive) connect with simplices that is not explained by differences in their in-and out-degrees. At this point it should be noted that the form of competitive inhibition explored in this work is different from lateral inhibition commonly described in the context of assemblies. Znamenskiy et al. (2024) and Lagzi et al. (2021) considered disjunct assemblies of neurons with relatively few or weak connections between them. In our case, competition is between groups that are both innervated by the same source neuron(s). Additionally, their assemblies were internally strongly wired, but without higher-order structure, while we focused specifically on the impact of such structure. It may be interesting to combine in the future the plasticity simulation setup of Lagzi et al. (2021) with assemblies that have a strong simplicial structure.

It should be noted that only a small fraction of neurons actually participated in simplices of the ”target” dimension we focused on (Fig. S11), leading to question whether our results have any significance for the rest of them. While not shown, our findings of excitatory divergence structuring inhibitory connectivity were also present when dimensions 4 or 5 were instead considered, albeit at lower strength. Participation in these structures should therefore be considered to be a matter of degrees, not simply true or false.

Our choice of using simplices is rooted in their previously predicted functional relevance (Reimann et al., 2017b; Nolte et al., 2020). Here, we have confirmed their functional relevance *in vivo*. Furthermore, the directed simplex was the second most overexpressed triad motif, implying strong structural relevance (Fig. S1). However, other motifs have also been studied, e.g., Song et al. (2005a), and Brunel (2016) count all motifs on 3 and up to 5 nodes respectively. The bound on the number of nodes in a motif is strongly driven by computational limitations. Indeed, a network on *N* nodes has *^N^* subnetworks on *n* nodes. In contrast, counting simplices can be done systematically across the size of the motif and has been implemented efficiently(Lütgehetmann et al., 2020). Indeed, for the central subnetwork of the MICrONS data set, counting all 2-simplices took 1.06 seconds contrasted with 72 minutes to approximate all triad motif counts. Even though cyclic motifs on *n*-nodes have similar generalization properties across dimensions as *n*-simplices, in practice simplex counts can be computed more efficiently, particularly in large, sparse graphs. This is the case despite the fact that theoretically they both have the same worst-case complexity (Johnson, 1975).

Given all this, we must ask: Does the higher-order structure have a purpose, or is it merely an epiphenomenon of connectivity? On the level of correlating structure with neuronal function, we have demonstrated a significant effect on pairwise spiking correlations in Fig. 1C. Similar roles for higher-order structure have been proposed before Reimann et al. (2017b); Nolte et al. (2020). Combining the MICrONS dataset with simulations of a biophysically detailed model and careful manipulations of its connectome, Egas Santander et al. (2024) have elevated this from correlations to showing causation. They found that the presence of high-dimensional simplices increases the reliability of spiking and hence the robustness of the cortical code, while participation of a neuron in fewer simplices enhances the efficiency of the code by reducing its correlation with other neurons. This not only demonstrates the relevance of the higher-order structure we uncovered, but also provides an explanation for the specific ”fan-out” structure observed: Neurons in the source position of a high-dimensional simplex participate in many simplices, that is, they are associated with subnetwork topologies that facilitate reliability. Conversely, neurons in the target position participate in fewer assemblies, that is, they are associated with topologies that improve coding efficiency. The reduced network in Fig. 5B, while a drastic simplification, can then be speculated to provide insights into the function of the circuit, after means to increase reliability and efficiency are stripped away.

We compared the results we found in the MICrONS data to a recently released, biologically detailed model of cortical circuitry (Reimann et al., 2024; Isbister et al., 2023). For virtually all analyses, we found the same result: The non-random trends are present in the nbS1 model, but weaker. The nbS1 model captures connectivity trends arising from the dendrite and axon shapes of neurons, along with their placement in the local brain geometry, and additional biological data such as synapses per connection distribution moments and bouton densities (Reimann et al., 2015). However, it lacks additional specificity, such as the strong preference of subclasses of VIP positive interneurons to innervate almost exclusively PV or SST positive neurons (Pi et al., 2013; Reimann et al., 2024). Additionally, connectivity in the model is instantiated randomly within the constraints and unshaped by activity dependent structural plasticity. Our results suggest that important organizational principles of neuronal connectivity are already present in such a naive state (i.e., unaffected by plasticity), and are further enhanced by plasticity. We implemented a heuristic algorithm to approximate the effects of structural plasticity to rewire excitatory connections consistent with results of a recent analysis of the MICrONS data (Ecker et al., 2023), and studied how it affected the connectivity trends. We found that it moved the model closer to the MICrONS data for all analysis focusing on the excitatory subgraph only, but further away for analyses of the structure of inhibitory relative to excitatory connectivity. This could indicate a failing of the rewiring algorithm, or alternatively that it needs to be paired with inhibitory plasticity. This, alongside with our overall results indicates strong specificity of inhibitory connectivity, shaped by structural rewiring of connections to and from inhibitory neurons in cortex. Our findings thus reveal an intricate interplay between structural plasticity of recurrent excitatory motifs, inhibitory mediated competition and disinhibition, and provide important constraints for the development of biologically constrained theories of neocortical plasticity and learning *in vivo*.

## METHODS

### Preparation of connectivity data

#### Electron microscopic dataset

We extracted the internal connectivity between 60,048 neurons in v117 of the “minnie65 public” release of the MICrONS dataset. Specifically we used the table‘’allen soma coarse cell class model v1minnie3 v1” to identify neurons and their types in broad classes. In that table, a small number of neurons were associated with duplicate entries in the column “pt root id”. We manually inspected three such examples in the MICrONS browser and found that they were apparently instances of two neurons being merged during meshing. To avoid possible errors, the rows corresponding to these neurons were removed from the table. Locations of neurons in the table were converted from voxel ids to nanometer based on the reported voxel resolution of 4 *×* 4 *×* 40*nm*.

For v1181 of the data instead the table “aibs metamodel mtypes v661 v2” was used and combined with the table “proofreading status and strategy” based on matching entries for “pt root id”. Missing entries in “proofreading status and strategy”, corresponding to non-proofread neurons, were filled in with the string “empty”. We refer to the result as the “neuron table”.

Information on synaptic connectivity, including synapse sizes were loaded from the table “synapses pni 2” in the version corresponding to the neuron table. Data were requested from MICrONS servers in chunks for 250 postsynaptic neurons at a time. In each chunk, synapses were matched to pre-and post-synaptic neurons based on the columns “pre pt root id” and “post pt root id”. We discarded rows where the presynaptic neuron was not found in our neuron table. After all chunks had been queried and processed, the results were concatenated into a single “synapse table”.

Finally, the neuron and synapse tables were wrapped into a data structure that we developed and optimized for connectomics and other graph-related analyses (“ConnectomeUtilities”; see Data and Code availability statement). The data structure support serialization to the hdf5 format and we have made the resulting file available.

#### Selection of neurons to analyze

We considered 15 centrally located excitatory subnetworks by filtering neurons according to their horizontal positions. They were admitted if they were inside a 500 *×* 300*µm* rectangle. For each subnetwork the rectangle was shifted by 50*µm* into another location, resulting in a 5 by 3 grid of subvolumes with significant overlap (Fig. 1A1). Only excitatory neurons were filtered according to these subvolumes; inhibitory neurons were considered for analyses independent of their location inside or outside the rectangle. When results for an individual subnetwork are presented, they are for the most central one. When data for all 15 subnetworks is shown, it provided as a mean and standard deviation taken over the subnetworks.

#### Distance-dependent controls

To assess significance, we compared results to distance-dependent control connectomes. For each subnetwork exponential control models were fit based on the observed distance-dependent connection probabilities within and between excitatory (E) and inhibitory (I) populations, i.e., one model each for E to E, E to I, I to E, and I to I. Models had the form 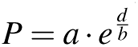, where *P* was the probability of connection, *d* was the Euclidean distance between neurons and *a* and *b* the fitted parameters. A stochastic instance was then generated, and analyses were conducted as in the baseline case.

#### Morphologically detailed model control

Additionally, we compared results to a recently released morphologically detailed model with connectivity derived from the shapes of neurons and biological constraints (Reimann et al., 2015; Isbister et al., 2023). This allows us to predict to what degree features of connectivity are explained by neuron morphologies, and to what degree other mechanisms, such as plasticity shape the connectome. For the model, subnetworks were only 300 *×* 300*µm*. The size was selected such that they contain approximately the same number of excitatory neurons as the MICrONS subvolumes. Inhibitory neurons were inside rectangles 100 *µm* larger in each of the four directions, yielding approximately the same number of inhibitory neurons.

### Topological analyses

#### Finding directed simplices and triad motifs

Using the connalysis (Egas Santander et al., 2024) python package (see Data and code availability), we calculated the list of directed simplices of all dimensions of the excitatory subnetwork. A directed simplex of dimension *dim* is a motif of *dim* + 1 neurons where the connectivity between them fulfills the following criterion: There exists a numbering of neurons, *i*, *j*, in the motif from 0 to *dim*, such that if *i < j* then there exists a synaptic connection from the *i*^th^ neuron to the *j*^th^ neuron (Fig. 1B). We call that numbering the position of the neuron in the simplex. We call neurons numbered near 0 the source side of a simplex and neurons near *dim* the target side. With the same package we computed estimates of triad motif counts of the central subnetwork of the MICrONS dataset. The estimate was obtained by classifying 5000, 000 connected triads that were randomly selected from the 152*^t^*981, 332 in the network and approximating the total values linearly.

#### Control model of simplex counts arising from reconstruction errors

We investigated a control model to explain the finding of elevated simplex counts in the central subvolume. The control network, a non-structured and undirected, i.e., Erdos-Renyi (ER), contains at least one simplex in the highest observed dimension, and is then subjected to loss of edges through reconstruction errors until the observed density of edges is reached. Kahle (2009) report that the expected maximal dimension in an undirected ER graph *G* on *n* nodes and connection probability *p* is *≈ −*2 log(*n*)*/* log(*p*). Note that these values are for undirected simplices and directed simplices are less likely to form because additionally the direction of each edge has to align. This estimate is therefore a conservative upper bound on the number of simplices to find in a directed ER graph. The central subnetwork we analyzed had 14559 nodes and maximal dimension 7. Based on the above formula, 13701468 edges would be required to form them in an undirected ER graph. To then reach the observed edge density of 819869 edges, 94% of edges would need to be removed. Therefore, the control can be rejected for reconstruction accuracies above 6% recall.

#### Analysis of inhibition of simplices

We then represented the topological location of all inhibitory neurons relative to the simplices as follows: Let *S_dim_* be the number of directed simplices in dimension *dim* and *N_inh_* be the number of inhibitory neurons. We constructed a *S_dim_ × dim* + 1 *× N_inh_* tensor *M_dim,ei_*, where the entry at index *i, j, k* is the number of synapses from the neuron at position *j* of excitatory simplex *i* to inhibitory neuron *k*. Similarly, in *M_dim,ie_* the entry at *i, j, k* is the number of synapses from the inhibitory neuron *k* to the neuron at position *j* of simplex *i*.

Let *Mʹ_dim,ei_* and *Mʹ_dim,ie_* be the sums of *M_dim,ei_* and *M_dim,ie_* over their second dimensions respectively. That is, *S_dim_ ×N_inh_* matrices that count the total number of synaptic connections from / to a simplex to / from an inhibitory neuron. Then we call the matrix product *Mʹ_dim,ei_* and *Mʹ⸆_dim,ie_* = *I_dim_* the matrix of disynaptic inhibition between simplices of dimension *dim*.

#### Symmetry of simplex-simplex inhibition matrices

We analyzed how symmetrical the matrix of inhibition between simplices was by calculating how much the variance of its entries is reduced when the transpose is subtracted:

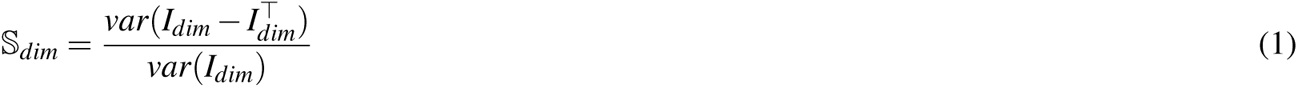

Increasing symmetry reduces this measure, until it reaches 0 in the case of perfect symmetry. The expected value if the entry at *i, j* and at *j, i* are statistically independent is 2. We calculated controls for *S_dim_* by shuffling the columns of *Mʹ_dim,ei_* and *Mʹ_dim,ie_* before calculating *I_dim_*and then *S_dim_* as usual.

#### Analysis of simplex overlap

Simplices can overlap in their constituent neurons by up to *dim* + 1 neurons. Let *O_dim,_*_[_ *_j,k_*_]_ be a *S_dim_ ×S_dim_* matrix that counts for each combination of simplices their overlap in neurons in positions between j and k (i.e. the size of the intersection of neurons in positions i with *j ≤ i ≤ k*).

#### Clustering of source-and target groups of simplices

Simplices were grouped using the Louvain clustering algorithm (Barber, 2007) from the python sknetwork package (Bonald et al., 2020) on the matrices *O*_6,[0,2]_ and *O*_6,[4,6]_ of the MICrONS subnetworks, yielding what we denote *source groups* and *target groups* of simplices respectively. The resolution parameter was set to 2.2.

#### Analysis of disinhibition

For inhibitory neurons, we calculated their *simplex indegree* as the number of connections from excitatory neurons in 6-dimensional simplices in positions 0, 1 or 2. Correspondingly, the *simplex outdegree* was the number of connections to neurons in 6-dimensional simplices in positions 4, 5 or 6. If a neuron participated in more than one simplex in one of the indicated positions its connection would be counted that number of times.

Also, for inhibitory neurons, we calculated their targeting preference for other inhibitory neurons as follows. We calculated for a neuron *i* its outdegree *d_i_^ttl^* and outdegree onto inhibitory neurons *d_i_^inh^*. We then compared against a control that keeps the total outdegrees of individual neurons and the overall fraction of connections onto inhibitory neurons. Under this null hypothesis, the probability to find an inhibitory outdegree as observed or higher is:

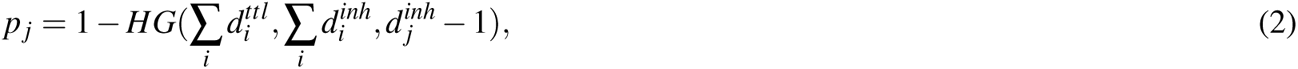

where *HG*(*M, N, n*) is the cumulative hypergeometric distribution of drawing *n* times from *M* objects with *N* positive objects. We considered a neuron to be inhibitory targeting if *p _j_ ≤* 10*^−^*^6^. The threshold was selected to yield around 15% of inhibitory neurons, in line with the fraction of inhibitory targeting cells in (Schneider-Mizell et al., 2023), which was 17%.

#### Neighborhood complexity

We measured the complexity of the graph neighborhoods of individual neurons in the MICrONS data in accordance with Egas Santander et al. (2024). Briefly, we first constructed as a control a configuration model of the central subnetwork, i.e., a random graph that preserves for each neuron its in-and out-degree. We then considered the subgraph given by the neighborhood of a neuron, i.e., the selected neuron and all its afferents and efferents and the connections between them. Specifically, we compared the neighborhood of a neuron with the neighborhood of the same neuron in the configuration model. As the control preserves degrees, both neighborhoods have the same size, but different structure. We compared them using the Wasserstein distance of the distribution of the degrees of their nodes.

#### Assessing effect of proofreading status and inhibitory subtypes

For the proofreading status of neurons we used version 1181 of the table “proofreading status and strategy” of the MICrONS data. To be able to analyze proofread and non-proofread neurons separately (or neurons that failed proofreading), we used slight variations of the analyses in Figs. 2-3.

While in 2 we consider the number of unique neurons in each simplex position, divided by the number of simplices, in Fig. 7C we instead divide by the number of neurons in each proofreading class. This was necessary for two reasons. First, it was individual neurons that were proofread, not simplices. The simplices were still detected in the network of both proofread and non-proofread neurons, as such a simplex could contain both types of neurons. Second, the number of proofread neurons was overall much lower than non-proofread ones. The analysis can still assess whether fewer unique neurons participate in one position than another.

While in Fig. 3D we depict the absolute inhibitory degree of neurons in different simplex positions, in Fig. 7D-F we additionally divide the results by the number of neurons in each proofreading class. This additional normalization yielded the connection probability with the indicated class. It was necessary because the number of proofread neurons was overall much lower than non-proofread ones.

Repeating the analysis for subtypes according to Schneider-Mizell et al. (2023) instead of proofreading class, allowed us to assess differences between these classes in how they interact with simplices. Note that while the inhbitory subtypes were first defined by Schneider-Mizell et al. (2023), the actual labels from the MICrONS data we used come from a soma and nucleus feature trained metamodel (Elabbady et al., 2022).

### Plasticity-inspired rewiring

We rewired the connectivity of the central subvolume of the nbS1 model with a custom algorithm inspired by recent findings, based on simulations and analysis of the MICrONS data (Ecker et al., 2023), showing that plasticity favors connections participating in many high-dimensional simplices. A single rewiring step worked as follows: First, for each excitatory connection (*a → b*) the maximum dimension of simplices it participates in was calculated as *D*(*a → b*). Next, for each excitatory neuron *a*, we counted the number of simplices it participated in, for each dimension and position and denote it by Par*_dim,pos_*(*a*). A specified number of connections *m* with a maximum dimension below or equal *D* were removed independently at random. Finally, the same number *m* new connections were added between random unconnected pairs of neurons, *a → b*, according to a probability, *P*(*a → b*). The probability was proportional to the participation of *a* and *b* in simplices above dimension *D*, in positions *i* and *j* respectively for *i < j* belonging to a specified set we denote pos as follows:

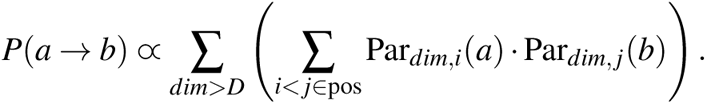

We conducted a single rewiring step with *D* = 3 (the middle dimension), *m* = 2000, followed by three steps with *D* = 2, *m* = 10000 each, for all we focused on the source side by using pos = *{*0, 1*}*.

### Analysis of functional MICrONS data

The MICrONS dataset we studied also contains functional data of neurons co-registered with the structural data. Specifically, it contains calcium imaging traces and spike trains derived from a deconvolution of the traces. The Pearson correlation of deconvolved spike trains of excitatory neurons was computed for 8 sessions. The sessions were selected from the activity recorded in the “minnie65 public” release version 661, such that at least 1000 neurons were scanned in each session and at least 85% of them are co-registered in the structural data we used. While every functionally imaged neuron is part of the EM volume, some were not contained in the table of neuron identifiers we used as the basis of the connectivity data (see above). Neurons that do not have a unique identifier were also filtered out (between 0 and 10.8% of co-registered cells and on average 2.8% across sessions). Next, we average these correlations along groups determined by the underlying structure in two ways. First, we grouped each excitatory connection by the maximum dimension of simplices it participates in. Second, for each dimension *n*, we extracted the list of the last connections in *n*-simplices (with repetitions), i.e., the connections between nodes in position *n−* 1 to *n*. Lastly, we averaged the correlations across those connections.

## ACKNOWLEDGMENTS

We thank Marwan Abdellah, Roberto Araya, Yamine Benabdesselam, Nghi Vuong Nguyen for helpful discussions.

This study was supported by funding to the Blue Brain Project, a research center of the Ècole polytechnique fédérale de Lausanne (EPFL), from the Swiss government’s ETH Board of the Swiss Federal Institutes of Technology.

E.B.M. was supported by funding from the Institute for Data Valorization (IVADO), the CHU Sainte-Justine Research Center (CHUSJRC), Fonds de Recherche du Québec–Santé (FRQS), the Canada CIFAR AI Chairs Program, the Quebec Institute for Artificial Intelligence (Mila), and Google. Compute infrastructure was supported in part through a grant of computing time to E.B.M. from the Digital Research Alliance of Canada.

## DATA AND CODE AVAILABILITY

- **Electron microscopic connectomics data**. We obtained the EM connectivity data from the pages of IARPA MICrONS (The MICrONS Consortium et al. (2021), https://www.microns-explorer. org/cortical-mm3).
- Version 117 of this data in a format that is optimized for the types of analyses presented here can be found under https://doi.org/10.5281/zenodo.8364070. It was used for the analyses in Figs. 1-6.
- Version 1181 of this data in the same format can be found under https://doi.org/10.5281/zenodo.13849415. It was used for the analyses in Fig. 7.
- **Connectivity of a morphologically detailed model**. We obtained the morphologically detailed model of (Reimann et al., 2024; Isbister et al., 2023) from their release under https://doi.org/10.5281/zenodo.8026353. A version of this data in a format that is optimized for the types of analyses presented here can be found under https://doi.org/10.5281/zenodo.10079406.
- **Loading the morphologically detailed model**. To load data from the morphologically detailed model, we used the bluepysnap python package (https://github.com/BlueBrain/ snap). It is published under https://doi.org/10.5281/zenodo.10090777.
- **Representation and basic analysis of connectomes**. Data loading, saving, filtering and ba-sic analyses were conducted using the ConnectomeUtilities python package (https://github.com/BlueBrain/ConnectomeUtilities). It is published under https://doi.org/10.5281/zenodo.10059227.
- **Topological analyses**. Detection of simplices and analyses related to them were conducted using the connalysis python package (https://github.com/danielaegassan/connectome_ analysis/). It is based on *flagsercount* (https://github.com/JasonPSmith/flagser-count).
- **Custom analysis code**. Plots were generated with custom python code, in jupyter notebooks. The code can be found under https://github.com/MWolfR/excitation_inhibition_topology. It requires all the python packages mentioned in the previous points.

## AUTHOR CONTRIBUTIONS

- Conceptualization: M.W.R, E.B.M
- Data curation: M.W.R.
- Formal analysis: M.W.R.
- Investigation: M.W.R.
- Methodology: M.W.R., D.E.S., A.E.
- Project administration: M.W.R.
- Software: M.W.R., D.E.S., A.E.
- Supervision: M.W.R., E.B.M.
- Validation: M.W.R., D.E.S.
- Visualization: M.W.R.
- Writing - original draft: M.W.R.
- Writing - review & editing: M.W.R., D.E.S., A.E., E.B.M.

## SUPPLEMENTARY INFORMATION

**Supplementary Figure 1.**
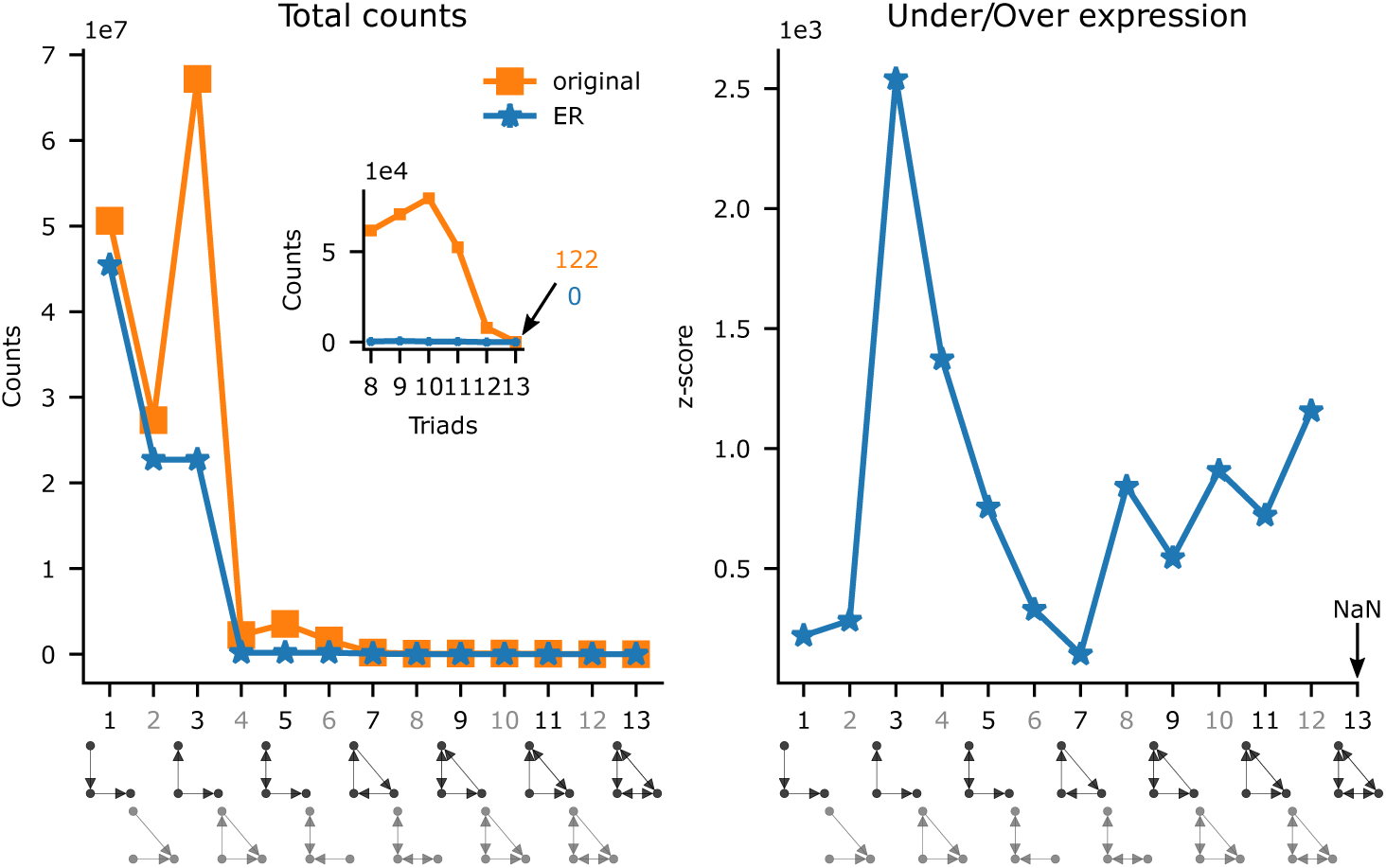
Left, triad motif counts. In blue, counts for the central subnetwork of the MICrONS dataset. In orange mean and std of the counts of 30 Erdős–Rényi controls. Right, MICrONS counts z-scored with respect to the distribution of the counts on the Erdős–Rényi controls.

**Supplementary Figure 2.**
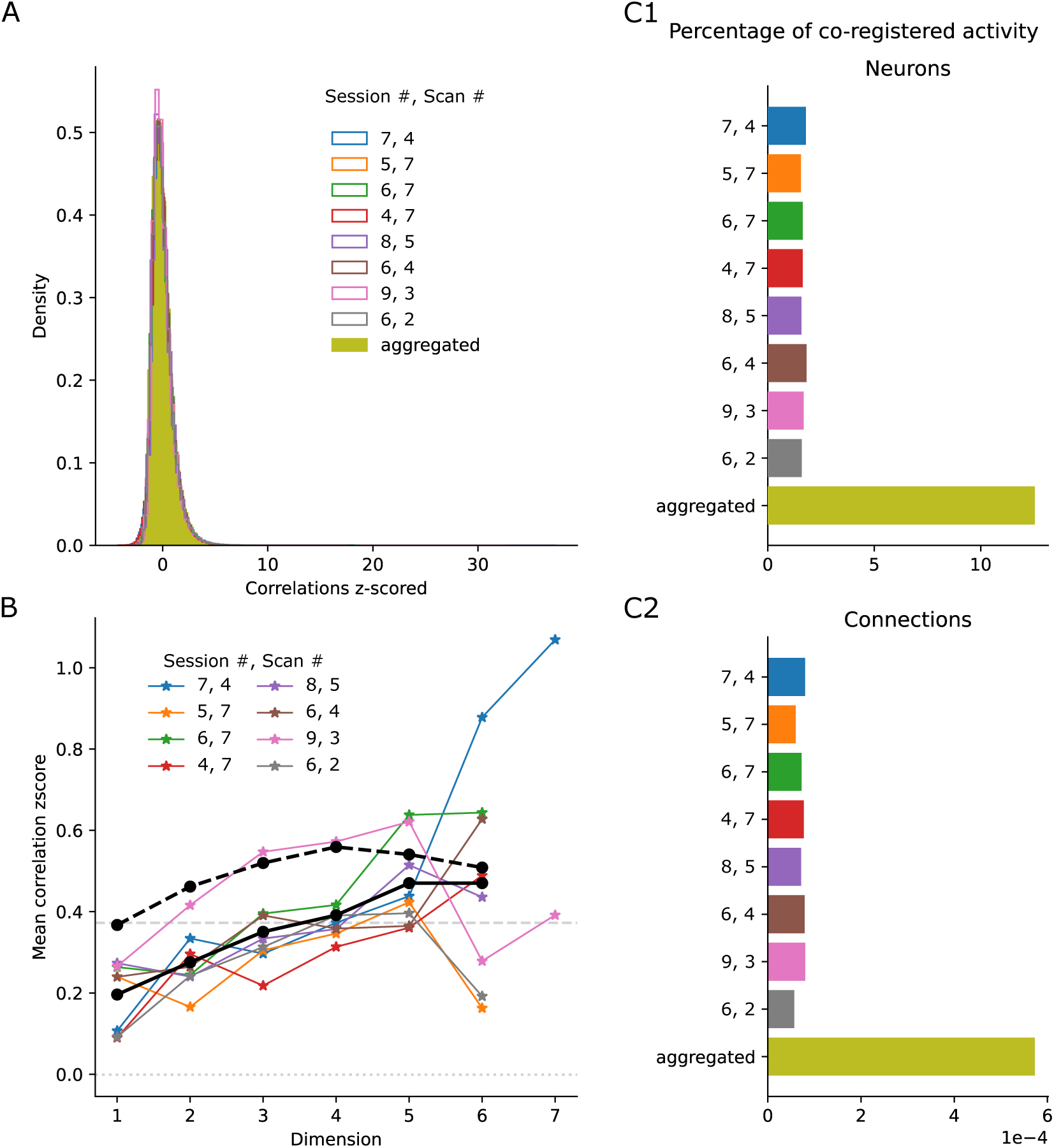
Activity per repetition in the MICrONS data set. A: Histograms of z-scored correlations for all repetitions. B1: Percentage of neurons with co-registered activity within the whole circuit in each repetition or aggregated across repetitions. B2: As B1 but for connections. C:Correlations of the activity (z-scored) of pairs of neurons against their simplex membership. The x-axis indicates the maximum dimension over simplices the connection participates in. Black lines: mean values over recording sessions. Black dashed line: values when only the last pair in a simplex is considered. Grey dashed line: Overall mean for connected pairs.

**Supplementary Figure 3.**
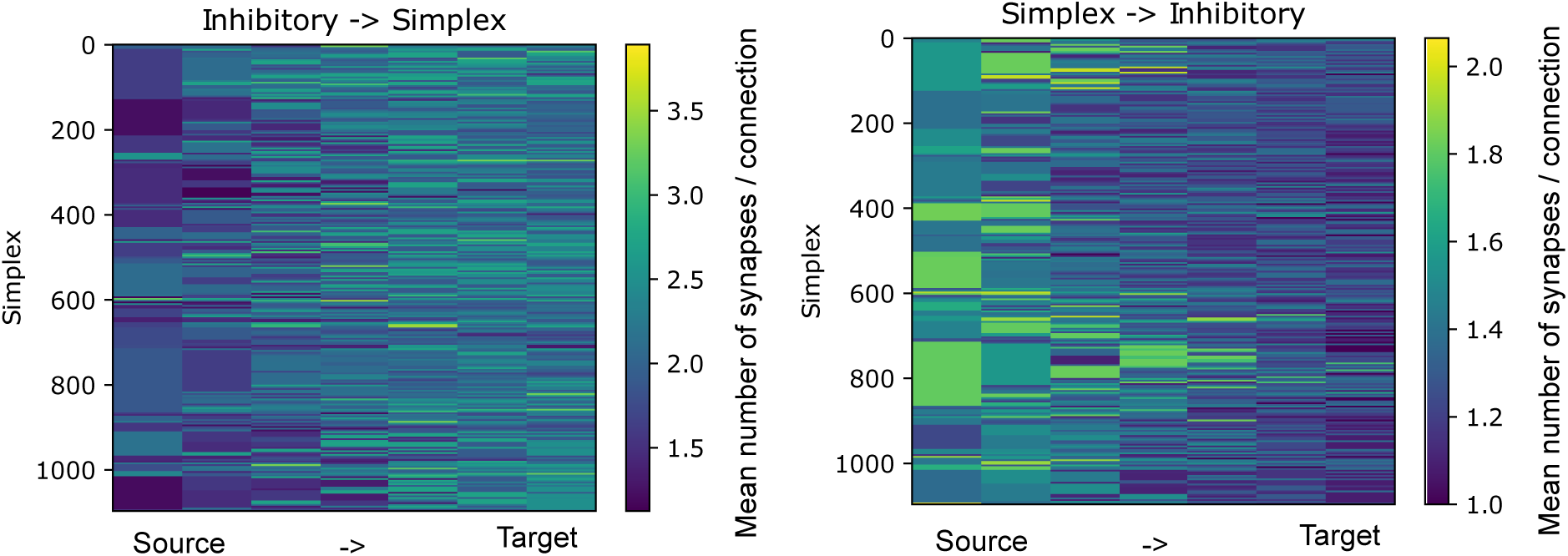
Inhibitory connectivity from / to 6-d simplices as in Fig. 3C, but instead of the degree (number of connections) we show the number of synapses per connection.

**Supplementary Figure 4.**
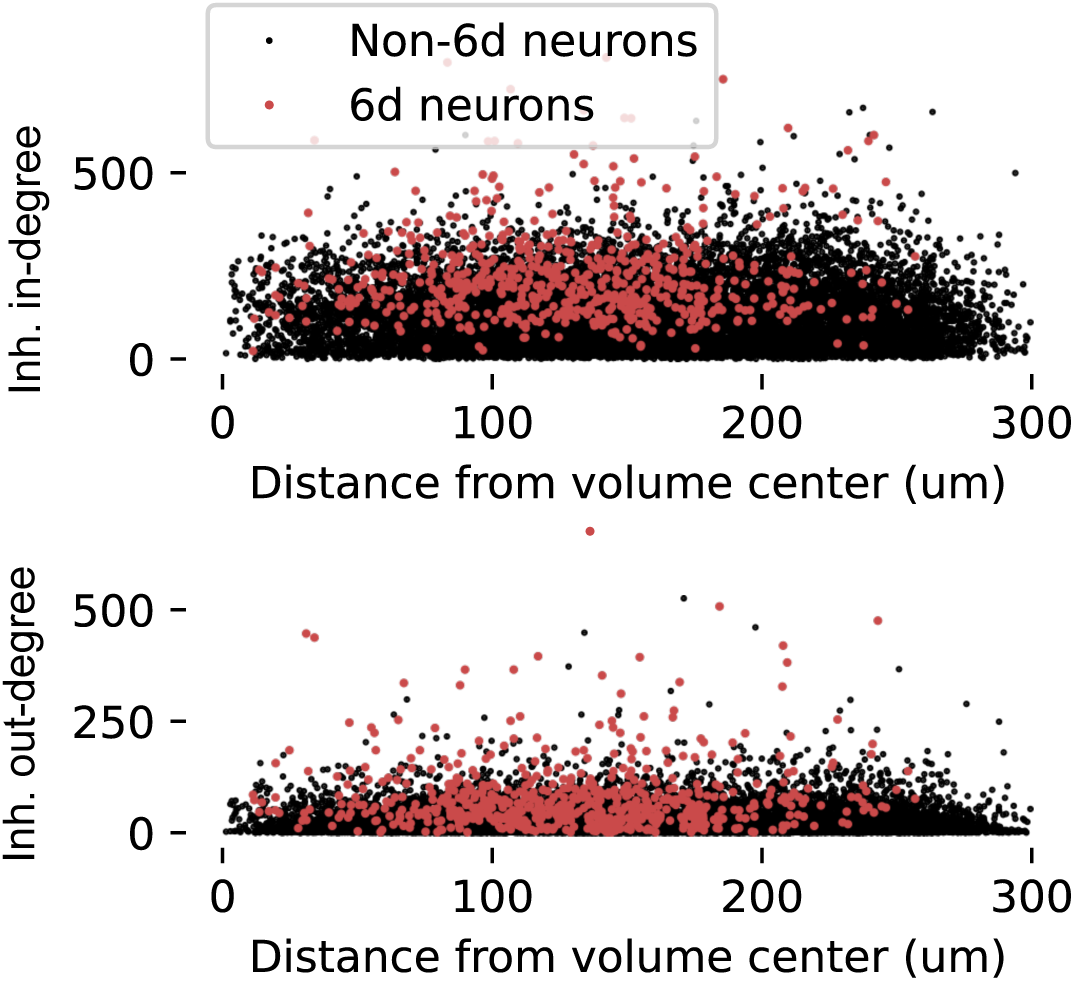
Distance from the volume center against inhibitory in-and out-degree for neurons in the central subnetwork of MICrONS. For members of the 6-core (red) and non-members (black).

**Supplementary Figure 5.**
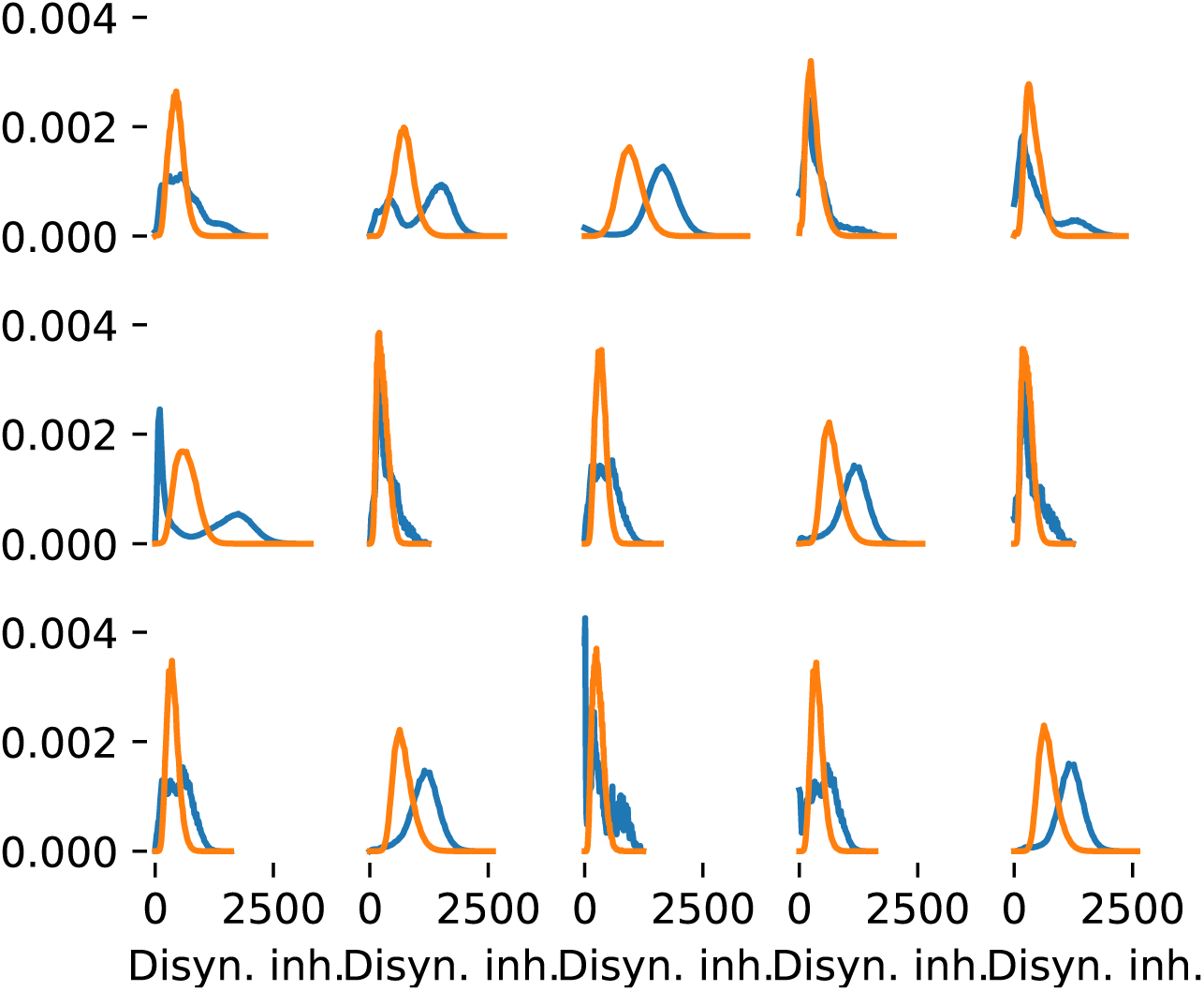
Disynaptic inhibition strength comparing the data (blue) against a control (orange) as in Fig. 4C for all subnetworks of MICrONS.

**Supplementary Figure 6.**
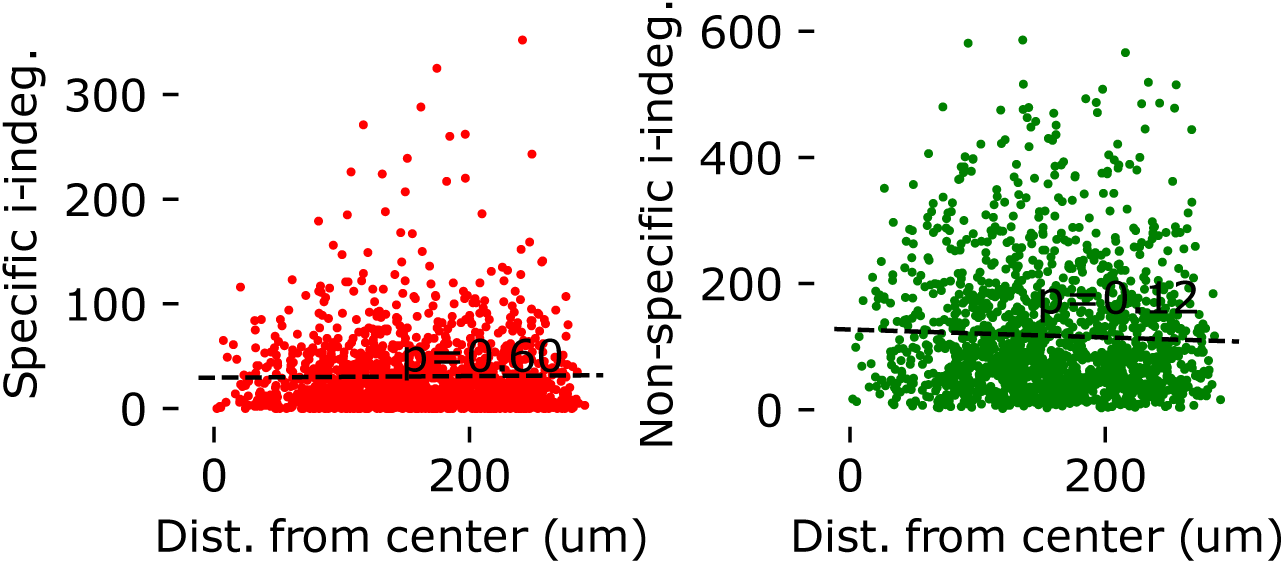
Distance from the volume center against inhibitory in-degree of inhibitory neurons in the central subnetwork of MICrONS. Left: from inhibitory neurons with a significant targeting preference for other inhibitory neurons. Correlation is non-significant (pearsonr: 0.013; p=0.60) Right: from neurons without or with weak targeting preference. Correlation also non-significant (pearsonr: -0.04; p=0.12).

**Supplementary Figure 7.**
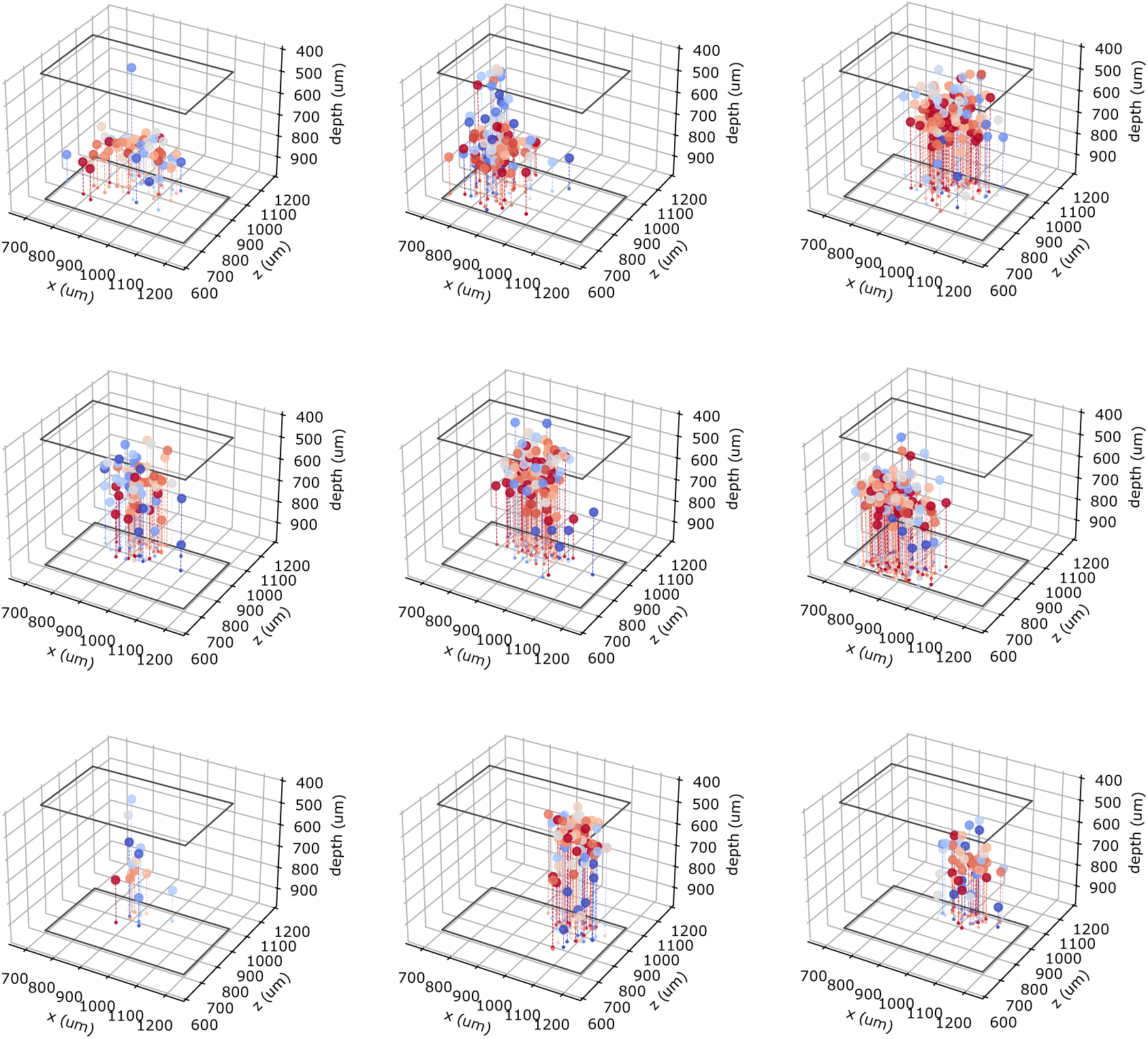
Spatial locations of neurons in source groups as in Fig. 6B. Groups 0-8 of Fig. 5B ordered from left to right, and top to bottom.

**Supplementary Figure 8.**
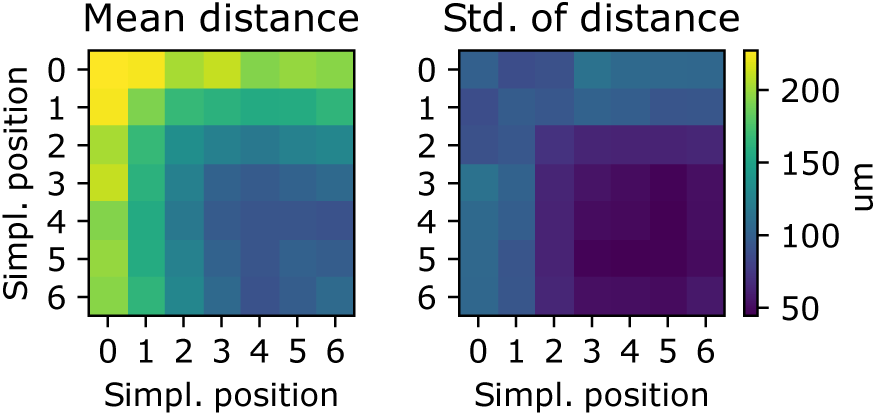
Pairwise distances of neurons in the same source group against their simplex positions. Given a pair of simplices in the same source group, we can calculate the distances between the neuron in position *i* of one simplex and the neuron in position *j* of the other, for all combinations of *i* and *j*. The plot shows the mean (left) and standard deviation (right) over all pairs of simplices in the same source group, for the source groups in Fig. S 7. Instances where neurons *i* and *j* were the same neuron, due to simplex overlap, were ignored.

**Supplementary Figure 9.**
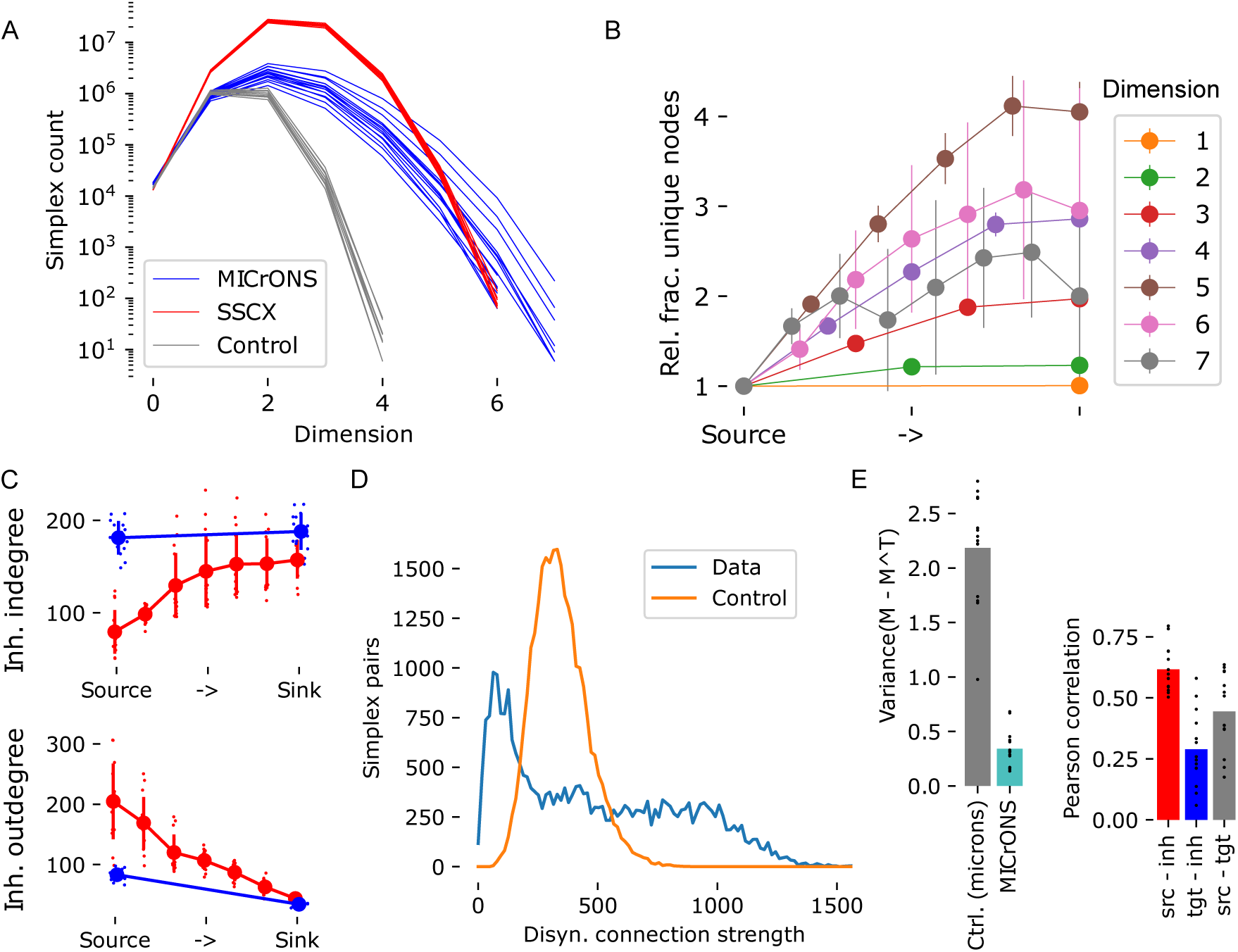
Repeating analyses for a later, more proofread version of the MICrONS dataset. A: As Fig. 1D. B: As Fig. 2C1. C: As Fig. 3D, top. D: As Fig. 4C, top. E, left: As Fig. 4D, left. E, right: As Fig. 4H, left.

**Supplementary Figure 10.**
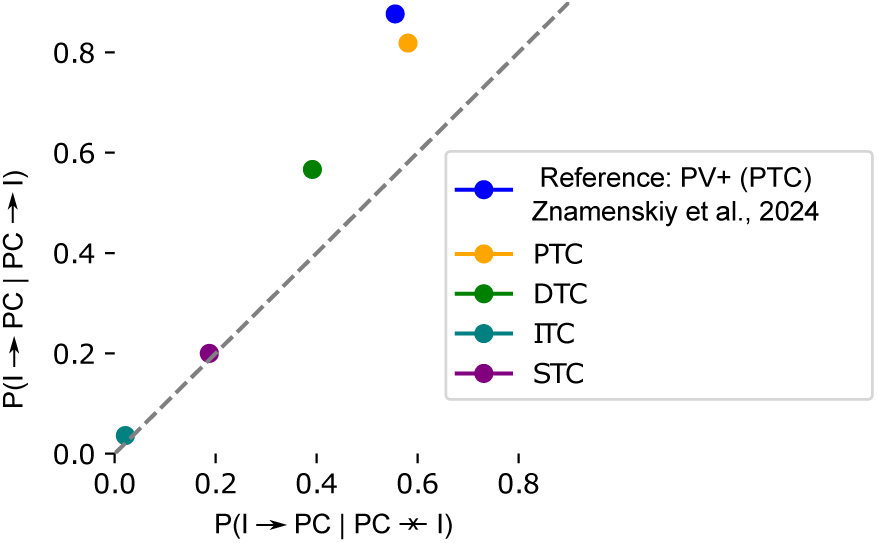
Comparing analyzed connectivity in MICrONS to recent experimental results. Znamenskiy et al. (2024) found an overexpression of reciprocal connectivity between PV-positive neurons and pyramidal cells in layer 2/3 of mouse visual cortex. We characterize the overexpression by contrasting the inhibitory to PC connection probability where the PC is not connected to the inhibitory neuron (x-axis) with the connection probability where the PC is connected (y-axis). This is conducted for the four classes of inhibitory neurons characterized by Schneider-Mizell et al. (2023); the “PTC” class corresponds tentatively to PV-positive neurons. The grey line indicates identity, i.e. no bias for or against reciprocal connectivity.

**Supplementary Figure 11.**
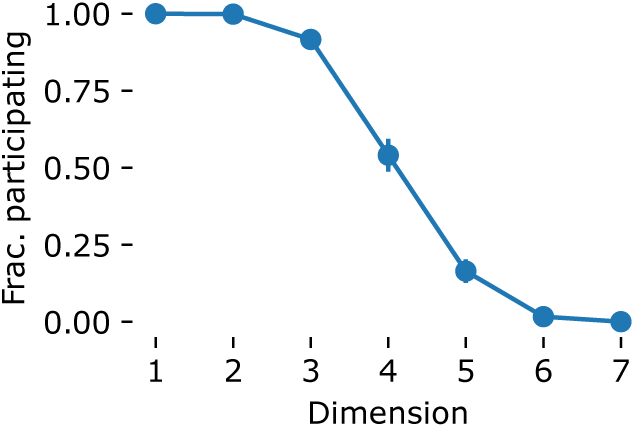
Fractions of excitatory neuron that participate in at least one simplex of the indicated dimensions.

